# Inflammation-Responsive Micellar Nanoparticles from Degradable Polyphosphoramidates for Targeted Delivery to Myocardial Infarction

**DOI:** 10.1101/2022.11.11.516224

**Authors:** Yifei Liang, Holly L. Sullivan, Kendal Carrow, Joanna Korpanty, Kendra Worthington, Colin Luo, Karen L. Christman, Nathan C. Gianneschi

## Abstract

Nanoparticles that undergo a localized morphology change to target areas of inflammation have been previously developed but are limited by their lack of biodegradability. In this paper, we describe a low ring strain cyclic olefin monomer, 1,3-dimethyl-2-phenoxy-1,3,4,7-tetrahydro-1,3,2-diazaphosphepine 2-oxide (MePTDO), that rapidly polymerizes via ring-opening metathesis polymerization (ROMP) at room temperature to generate well-defined degradable polyphosphoramidates with high monomer conversion (>84%). Efficient MePTDO copolymerizations with norbornene-based monomers are demonstrated, including a norbornenyl monomer functionalized with a peptide substrate for inflammation-associated matrix metalloproteinases (MMPs). The resulting amphiphilic peptide brush copolymers self-assembled in aqueous solution to generate micellar nanoparticles (30 nm in diameter) which exhibit excellent cyto- and hemocompatibility and undergo MMP-induced assembly into micron scale aggregates. As MMPs are upregulated in the heart post-myocardial infarction (MI), the MMP-responsive micelles were applied to target and accumulate in the infarcted heart following intravenous administration in a rat model of MI. These particles displayed a distinct biodistribution and clearance pattern in comparison to non-degradable analogues. Specifically, accumulation at the site of MI, competed with elimination predominantly through the kidney rather than the liver. Together, these results suggest this as a promising new biodegradable platform for inflammation targeted delivery.

## Introduction

Matrix metalloproteinases (MMPs) are upregulated in inflammatory environments, including in the heart following acute myocardial infarction (MI). Previously, we reported that MMP-responsive micellar nanoparticles, assembled from MMP-responsive peptide-functionalized polynorbornene amphiphiles, localize efficiently to the infarcted heart following systematic administration in a rat MI model.^1^ This occurs when the initially small diameter, spherical micelles (20 nm in diameter) extravasate, enter the muscle tissue, and interact with MMPs which induce via a phase change from spheres to micron-scale assemblies upon depletion of the polar head group by MMP enzymes. However, these materials were not biodegradable, limiting translational potential. Indeed, degradable nanoparticles have been employed in delivery to MI in the past but have not been shown to efficiently accumulate within the disease site for an extended period of time.^2–4^ Herein, we aim to combine enzyme responsiveness for targeting and degradable polymer backbones to respectively accumulate and clear materials on relevant timescales.

In recent years, ring-opening metathesis polymerization (ROMP), which generates well-defined polymers with diverse structure and function,^5–9^ has been further explored in the context of generating polymers with degradable backbones.^10^ Given the excellent functional group tolerance, monomers with a variety of degradable linkages, such as acetal/ketal,^11–13^ carbonate,^14^ oxazinone,^15^ enol ether,^16^ silyl ether,^17^ phosphoester,^18^ and phosphoramidate,^19^ have been successfully polymerized via ROMP. With judicious selection of polymerization conditions, such as catalyst, monomer concentrations, reaction temperature and solvent, degradable polymers with targeted molecular weights and narrow molar mass distribution can be achieved.^20–22^ Lately, functional degradable systems have been described in the context of biologically active materials. In 2019, Kiessling and coworkers demonstrated that fluorophores, reporter groups and bioactive epitopes can be introduced to ROMP-derived polyoxazinones through postpolymerization modification using oxime chemistry.^23^ In 2020, Johnson and coworkers described the copolymerization of silyl ether monomers with a PEGylated macromonomer and demonstrated the *in vivo* biodistribution and clearance of the resulting materials.^17^ Given these rare, yet promising examples, new monomers with high ROMP reactivity and facile functionalization are desirable. To this end, we previously reported the synthesis of well-defined and fully degradable polyphosphoramidates via low temperature ROMP of a low ring strain diazaphosphepine based cyclic olefin, 2-phenoxy-1,3,4,7-tetrahydro-1,3,2-diazaphosphepine 2-oxide (PTDO, **Figure 1**).^19^ Despite this initial success, some fundamental drawbacks hindered its further development. First, PTDO has limited solubility (0.3 M at 20 °C) in commonly used ROMP solvents such as dichloromethane (DCM) and tetrahydrofuran (THF). Second, low monomer conversion (55% under optimal conditions) limits scalability and general utility.

**Figure 1.**
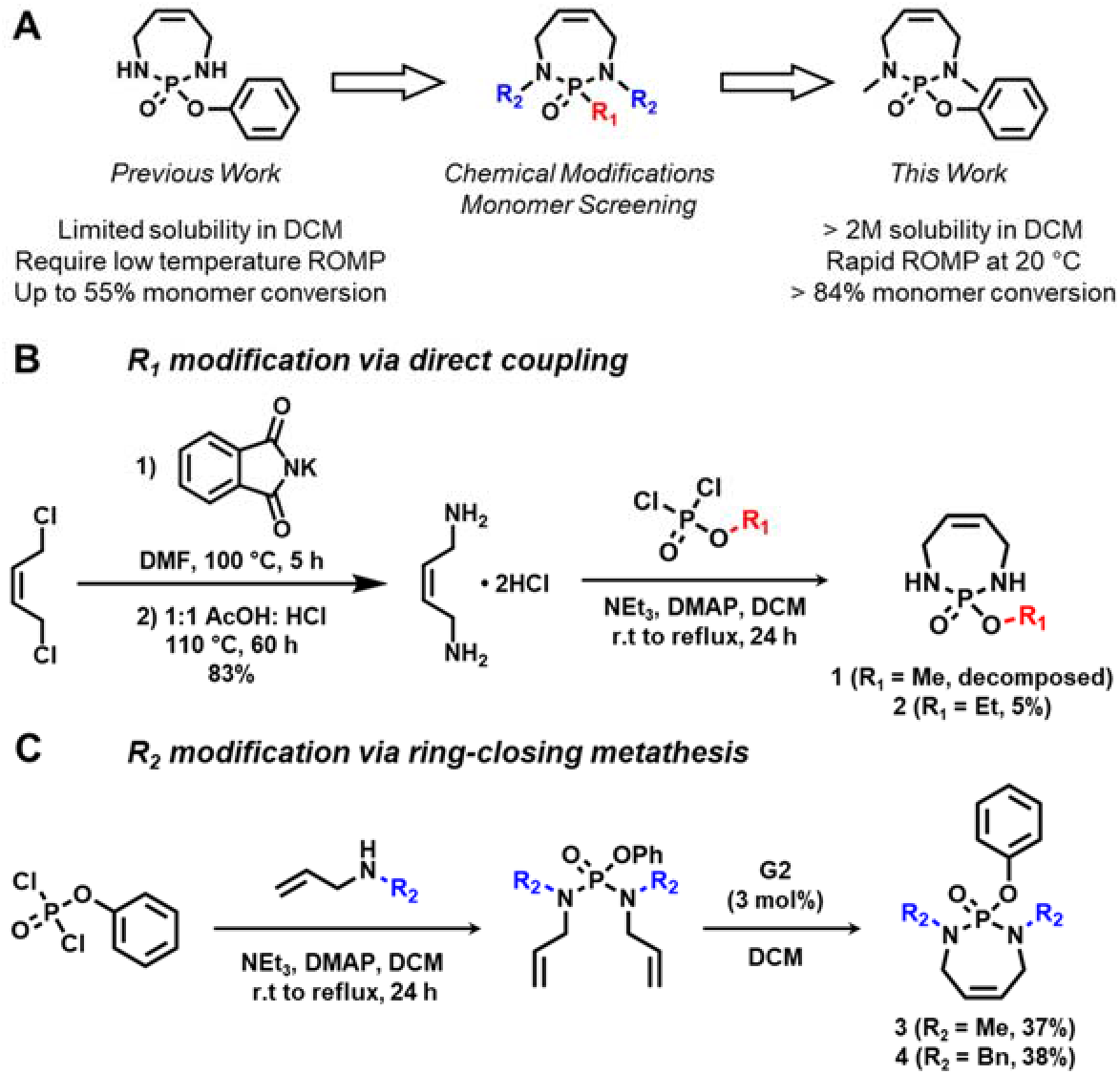
Monomer screening and synthesis of phosphoramidate monomers. (A) Development of MePTDO by using PTDO as the starting point. (B) Synthesis of R_1_ modified phosphoramidate monomers through direct coupling approach. (C) Synthesis of R_2_ modified phosphoramidate monomers through ring-closing metathesis.

Herein, we report a new phosphoramidate containing cyclic olefin, 1,3-dimethyl-2-phenoxy-1,3,4,7-tetrahydro-1,3,2-diazaphosphepine 2-oxide (MePTDO), which can be dissolved up to 2 M in DCM and has superior ROMP reactivity as compared to PTDO (**Figure 1A**). At room temperature, greater than 84% MePTDO conversion was achieved within 30 min in DCM, giving rise to high molecular weight (up to 80 kDa) polyphosphoramidates with narrow dispersity and backbone degradability. Furthermore, MePTDO exhibited similar ROMP reactivity in DMF, which is uncommon for low strain cyclic olefins. MePTDO also efficiently copolymerized with functionalized norbornenes, including a norbornenyl-peptide monomer containing a substrate for inflammation-associated matrix metalloproteinases (MMPs). We demonstrate that degradable peptide brushes based polyphosphoramidate amphiphiles can assemble into spherical micelles and respond to MMPs via cleavage of peptides and subsequent phase transition. This inflammatory enzyme responsiveness was subsequently leveraged for *in vivo* targeting. We demonstrated the degradable micellar nanoparticles (DNPs) can selectively accumulate in the infarcted heart in a rat acute MI model and displayed a distinct biodistribution and clearance pattern predominantly through the kidney.

## Results and Discussion

### Monomer Synthesis and Screening

A small library of phosphoramidate monomers was prepared using PTDO as the starting point. Substituents were introduced either at the phosphorous center (R_1_) or to the secondary amine (R_2_) with the goal of modulating ROMP reactivity (**Figure 1**). We hypothesized that sterically large functional groups at R_1_ would stabilize the ring-closed form due to the Thorpe-Ingold effect, increasing the equilibrium monomer concentration required for polymerization to occur.^24–26^ By replacing the phenoxy group with a smaller methoxy or ethoxy group, we aimed to destabilize the ring structure favoring ring opening and subsequent polymerization. On the other hand, a change from secondary to tertiary amine by introducing R_2_ would reduce unwanted amine-initiator coordination, thereby facilitating polymer chain growth and extending catalyst lifetime.^27–29^

To begin our investigation, R_1_ substituted monomers were approached by reacting cis-1,4-diamino-2-butene·2HCl with different dichlorophosphates (**Figure 1B**). However, we were not able to isolate monomer **1** due to the presence of side reactions and monomer decomposition. Similarly, only a trace amount (5%) of **2** was isolated from the coupling reaction. These results indicated that the phenoxy group is critical in maintaining the structural integrity of the ring and should not be replaced. To prepare R_2_ modified monomers, we took a ring-closing metathesis (RCM) approach (**Figure 1C**).^30^ Dienes were first synthesized by coupling dichlorophosphate with alkylated allylamine and cyclized via RCM in the presence of 3% 2^nd^ generation Grubbs catalyst **G2**. **G2** was subsequently removed by overnight absorption with basic aluminum oxide followed by column chromatography,^31^ yielding both **3** and **4** as white solids. Inductively coupled plasma mass spectrometry (ICP-MS) analysis revealed 0.54 ppm of residual Ru in **3**, which is negligible and should not affect the subsequent ROMP reaction.

To estimate the ring strain energy (RSE) of monomers **2** to **4**, density functional theory (DFT) calculations were performed. As shown in **Table S1**, all modified monomers have a lower RSE as compared to PTDO (10.86 kcal/mol). In particular, we note a negative RSE for monomer **4**, suggesting the ring-closed structure is more stable than the ring-opened form. This could be caused by the Thorpe–Ingold effect as the large benzyl groups forced ring closure. To examine the accuracy of the calculated RSE, a similar DFT method was used to estimate the RSE of norbornene. As compared to the experimental value (27.2 kcal/mol), the 16.2 kcal/mol from DFT is markedly lower.^32^ This discrepancy indicates DFT calculations may underestimate the RSE of these phosphoramidate monomers. Therefore, test polymerizations were performed using the previously identified low temperature ROMP conditions (2 °C, [M]_0_ = 0.5 M, 5 h in 10/90 v/v MeOH/DCM).^19^ Among all three monomers, only **3** successfully polymerized, giving rise to a polymer with expected *M*_n_ = 7000 and *Ð* = 1.08 (**Figure S1**). Despite nearly 50% monomer **2** conversion determined by ^1^H nuclear magnetic resonance (NMR), only oligomers were observed via size-exclusion chromatography coupled with multiangle light scattering (SEC-MALS), indicating significant backbiting. This result is unsurprising given the low RSE. In addition, the small substituent group (i.e., ethyl) may not provide enough steric shielding, thereby increasing the risk of secondary metathesis and decomposition of the propagating carbene.^33, 34^ As for **4**, no reaction was observed either through ^1^H NMR or SEC-MALS analysis, which correlates well with the negative RSE. Giving these results, in-depth investigations were performed on **3**, which is subsequently referred to as MePTDO in the following discussion.

### MePTDO Homopolymerization

Unlike PTDO, MePTDO is well solubilized in pure DCM (> 2M) and can polymerize at room temperature (20 °C). No significant difference in monomer conversion or polymer molar mass was noticed as compared to the reaction at 2 °C (**Figure S1**). Because of this, parallel reactions were set up in DCM at 20 °C by adding 0.5 M MePTDO to 0.01 equiv. of 3^rd^ generation Grubbs’ catalyst **G3**. The reactions were terminated at different times with ethyl vinyl ether (EVE) to investigate the polymerization kinetics. The polymerization reached equilibrium within 30 min with ~70% monomer conversion, affording a well-defined polymer with expected molecular weight and narrow dispersity (*M*_n_ = 24.3 kDa, *Ð* = 1.17) (**Figure S2** and **Table 1** Entries 1-4). No significant change in *M*_n_ or *Ð* was observed after 24 h, suggesting minimal chain transfer of backbone olefins. Since MePTDO has a lower RSE as compared to the original PTDO, we attributed this fast polymerization kinetics to the reduction in amine-initiator interactions. The steric hindrance caused by methylation potentially prevents the amine from coordinating the propagating Ru carbene, and thus allows the incoming olefin to properly coordinate for ring opening (**Figure S3**). Furthermore, the methyl group may provide shielding on the nearby propagating carbene, thereby reducing secondary metathesis and catalyst decomposition.^33–35^

**Table 1.** Scope of polymerization conditions ^*a*^

| Entry | [M]<sub>0</sub> (M) | [M]<sub>0</sub>: [**G3**] | Time (h) | Conv. <sup>b</sup> | $M_{n, \text{theo}}$<br>(KDa) <sup>c</sup> | $M_{n, \text{MALS}}$<br>(KDa) <sup>d</sup> | $\bar{D}$ <sup>d</sup> |
| --- | --- | --- | --- | --- | --- | --- | --- |
| 1 | 0.5 | 100:1 | 0.5 | 71 | 17.9 | 24.3 | 1.17 |
| 2 | 0.5 | 100:1 | 1 | 73 | 18.4 | 25.0 | 1.19 |
| 3 | 0.5 | 100:1 | 3 | 73 | 18.4 | 26.6 | 1.18 |
| 4 | 0.5 | 100:1 | 24 | 75 | 18.9 | 25.6 | 1.18 |
| 5 | 0.1 | 100:1 | 24 | No detectable polymer |  |  |  |
| 6 | 1.0 | 100:1 | 1 | 88 | 22.2 | 31.4 | 1.20 |
| 7 | 2.0 | 100:1 | 1 | 94 | 23.7 | 35.5 | 1.22 |
| 8 | 1.0 | 200:1 | 1 | 86 | 43.4 | 46.6 | 1.20 |
| 9 | 1.0 | 500:1 | 1 | 70 | 88.3 | 80.8 | 1.24 |
| 10 <sup>e</sup> | 1.0 | 100:1 | 1 | 84 | 21.2 | 26.8 | 1.18 |

<sup>a</sup>ROMP was performed in DCM under N<sub>2</sub> at room temperature and chain terminated with ethyl vinyl ether (EVE). <sup>b</sup>Monomer conversion was determined by $^1\text{H}$ NMR spectroscopy. <sup>c</sup>Theoretical molecular weight was calculated by $M_{n, \text{theo}} = \sum \text{DP}_{\text{monomer}} \times \text{MW}_{\text{monomer}}$ . <sup>d</sup>Molecular weight and dispersity were determined by SEC-MALS with a dn/dc of 0.179 mL/g. <sup>e</sup>Polymerization was performed in DMF.

Next, we investigated the impact of initial monomer concentration [M]_0_ on the polymerization. When [M]_0_ of 0.1 M was used, no reaction was observed after 24 h, suggesting 0.1 M is below the equilibrium monomer concentration required to initiate the polymerization.^10, 21^ Since MePTDO is well solubilized in pure DCM, we were able to perform the reactions with [M]_0_ of 1.0 and 2.0 M. Indeed, the increase in [M]_0_ enhanced monomer conversion to 94%, which is significantly higher than the 55% monomer conversion for the original PTDO (**Figure S2**, **Table 1** Entries 5-7).^19^ Further, by changing the monomer to catalyst feed ratio, P(MePTDO) with different degrees of polymerization (DPs) and narrow dispersity were obtained (**Figure S2**, **Table 1** Entries 6, 8-9). The measured molecular weights closely matched the theoretical values, indicating a living-like polymerization. Nevertheless, when the ratio of MePTDO:**G3** was increased to 500:1, significant gelation was observed, which led to a decrease in monomer conversion. Subsequently, we found MePTDO can efficiently polymerize in DMF, which is typically considered a poor solvent for ROMP of low strain cyclic olefins. 84% monomer conversion was achieved in 1 h when 1.0 M MePTDO was used (**Figure S2** and **Table 1** Entry 10). This improved solvent compatibility enabled us to investigate the copolymerization of MePTDO with peptide-functionalized norbornenes (*vide infra*).

### Acid Catalyzed Hydrolysis of MePTDO and P(MePTDO)

To study the degradation of the modified phosphoramidate systems, both MePTDO and P(MePTDO) (**Table 1** Entry 6) were subjected to 0.25 M DCl in DMSO-*d6*. The degradation was monitored by ^1^H and ^31^P NMR spectroscopy, as well as SEC-MALS (**Figure 2**, **Figure S4-5**). As shown by the NMR spectra (**Figure S4**), MePTDO rapidly hydrolyzed into phosphoric acid within 24 h. By integrating the residual MePTDO alkene signals with respective to the aromatic proton signals, a half-life (t1/2) of 6.5 h was revealed (**Figure 2A**). The peak at 4.0 ppm on ^31^P NMR was attributed to the partially hydrolyzed intermediate species (**Figure S4**). The low intensity suggested a fast turnover from the intermediate to the final degradation products. This degradation pattern is distinct from that for PTDO, where 1:1 intermediate: phosphoric acid was detected at 48 h post acid treatment.^19^ This discrepancy is attributed to the different pKa of leaving amine and agrees well with the findings from Berkman and coworkers who reported that their phosphoramidate scaffolds with tertiary amines hydrolyzed faster than the secondary amine analogs.^36, 37^ For P(MePTDO), NMR analysis revealed > 90% polymer degradation in 280 h, giving a t1/2 of 110 h (**Figure 2B**, **Figure S5**). The decrease in polymer NMR signal correlates well with the polymer molar mass reduction on SEC-MALS (**Figure 2C**). Like the monomer, only trace intermediate species was observed during polymer degradation (**Figure S5**).

**Figure 2.**
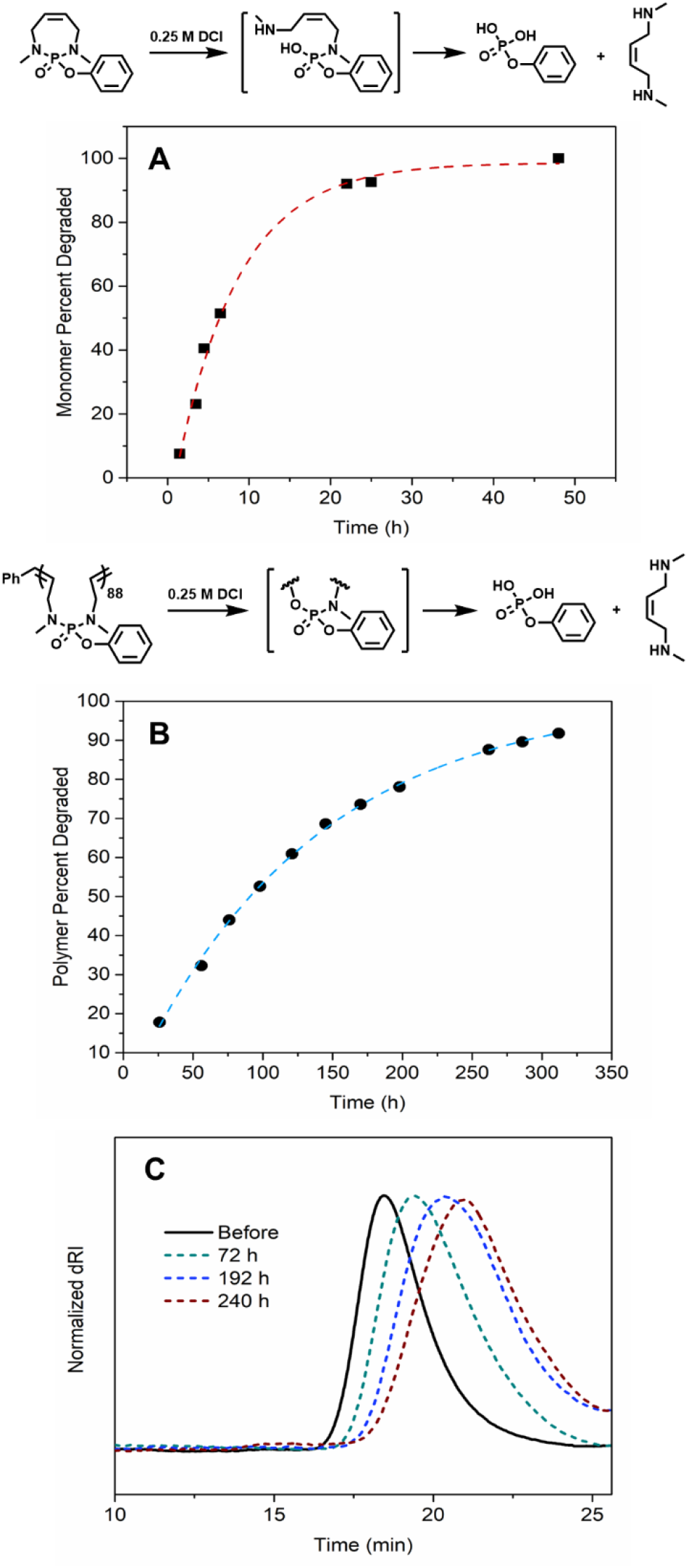
Degradation of MePTDO and P(MePTDO) monitored via NMR and SEC-MALS. (A) Percent MePTDO degradation versus time determined by ^1^H NMR. (B) Percent P(MePTDO) degradation versus time determined by ^1^H NMR. (C) SEC traces of P(MePTDO) degradation at different times.

### Physical Properties of P(MePTDO)

At room temperature, P(MePTDO) (**Table 1** Entry 6) is found to be a soft, rubbery solid. A glass transition temperature (T_g_) of −3 °C was determined by differential scanning calorimetry (DSC) (**Figure S6**). This T_g_ is markedly lower than the 42 °C of P(PTDO), potentially due to the reduction of interchain attractions and cohesion attributed to the methylation. This feature is attractive for responsive systems where phase transitions directed by changes in polymer head-group geometry drive changes in packing in resulting micelles.

### Scope of Norbornene Comonomers and Synthesis of Peptide Brush Polyphosphoramidate

To explore whether MePTDO would copolymerize with other ROMP monomers, we performed test reactions with norbornenes bearing different functional groups, including phenyl (NorPh), N-hydroxysuccinimide (NorNHS), and a peptide substrate of inflammation associated matrix metalloproteinase (NorMMP) (**Figure 3A**, **Figure S7-9**). For copolymerization with NorPh and NorNHS, 50:50:1 MePTDO: norbornene: **G3** was used and [MePTDO]_0_ was kept at 0.5 M in DCM (**Figure S7A-B**). To form random copolymers, norbornene and MePTDO were mixed before addition to **G3**. The reaction was terminated with EVE after 1.5 h and the polymer was precipitated by addition of cold diethyl ether. When NorPh was used as the comonomer, ^1^H NMR analysis revealed 100% NorPh consumption and 70% MePTDO incorporation, giving rise to NorPh_50_-*co*-MePTDO_35_ (*Ð* = 1.24). As for the reaction with NorNHS, despite complete NorNHS consumption, only 30% MePTDO was converted, affording NorNHS_50_-*co*-MePTDO_15_ (*Ð* = 1.25). This limited conversion is possibly due to polymerization-induced gelation, which prevents further MePTDO incorporation. Both copolymers were treated with 0.5 M HCl in DMF for 48 h to achieve accelerated degradation. The shift and broadening of SEC signals confirmed polymer degradation into oligomers, suggesting MePTDO was incorporated in a stochastic manner in the polyolefin backbone (**Figure S7C-D**). To prepare block copolymers, norbornene was allowed to react with **G3** for 30 min before MePTDO addition. Although NorPh_50_-*b*-MePTDO_40_ (*Ð* = 1.11) was successfully synthesized, significant gelation was observed post NorNHS block formation, preventing chain extension. This observation suggested that the norbornene imide type monomer is more compatible with MePTDO. Upon acid treatment on NorPh_50_-*b*-MePTDO_40_, the MePTDO block was completely degraded, giving rise to two SEC peaks: one from the nondegradable polynorbornene block (18 min) and the other from small molecule residuals post polyphosphoramidate degradation (20~25 min) (**Figure S7E**).

**Figure 3.**
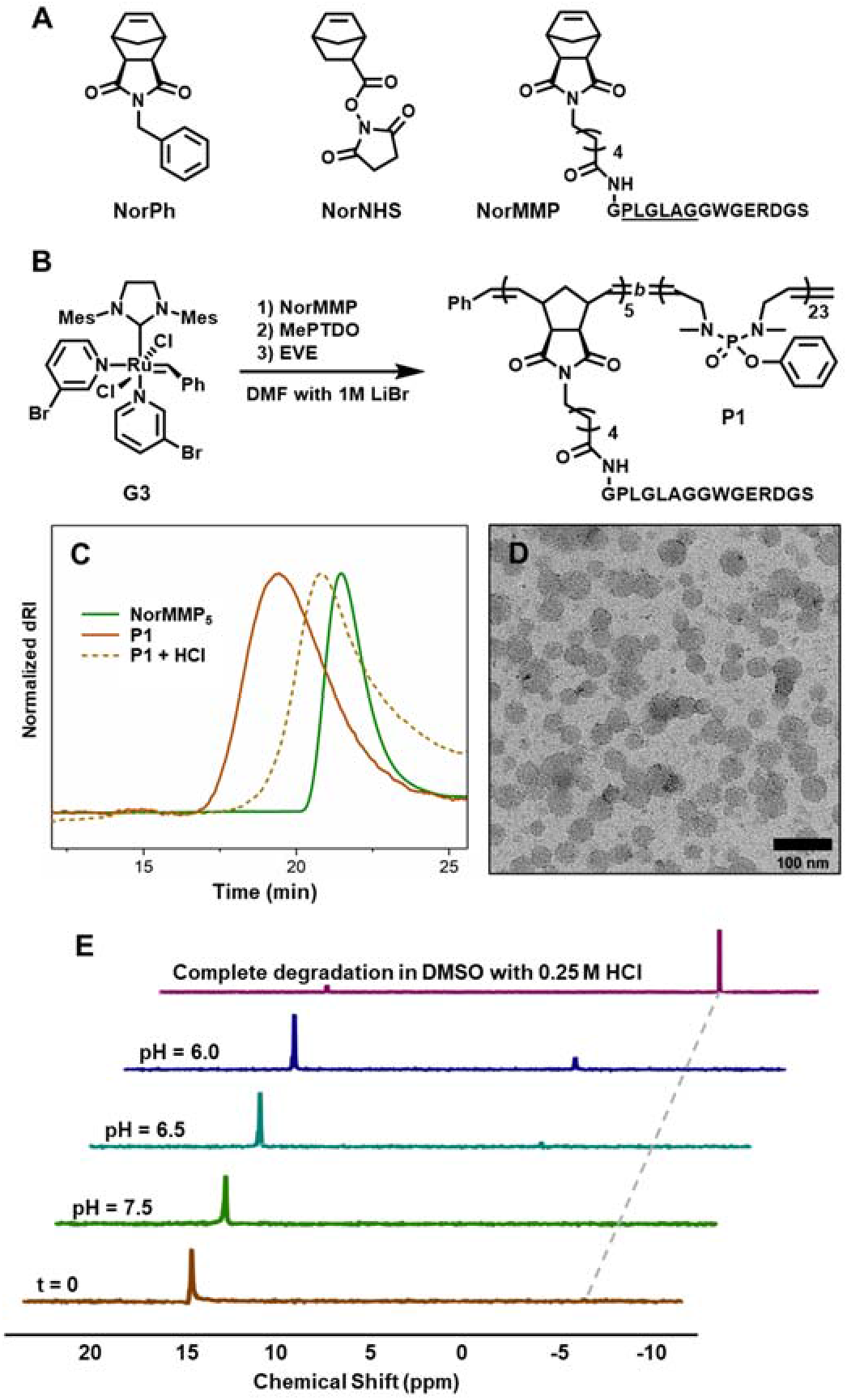
Scope of norbornene comonomers and the stability of nanoparticles formulated from peptide brush polyphosphoramidate **P1**. (A) Structures of norbornene comonomers. (B) Graft-through ROMP for the synthesis of peptide brush polyphosphoramidate **P1**. (C) SEC traces of **P1** before and after accelerated degradation in 0.5 M HCl in DMF. NorMMP_5_, which was generated by removing an aliquot of reaction mixture before MePTDO addition and terminated with EVE, was analyzed to confirm the step growth of polymer molar mass. (D) TEM image of **P1** assembly into spherical micelles. (E) ^31^P NMR spectra of **P1** after 10 days in DPBS buffer at pH of 6.0, 6.5 and 7.5. Spectra at t = 0 and post accelerated degradation in DMSO were included as reference. The dotted line at −6.4 ppm indicates the phosphoric acid peak.

To further demonstrate the versatility of this phosphoramidate system, a peptide brush polyphosphoramidate **P1** was prepared via graft-through ROMP (**Figure 3B**). NorMMP was allowed to completely polymerize before the addition of MePTDO (75% conversion).^1, 38, 39^ Upon acid treatment, the polyphosphoramidate block of **P1** was selectively degraded (**Figure 3C**). By dialyzing from DMSO into DPBS, **P1** assembled into spherical micelles with 20~50 nm in diameter as visualized by transmission electron microscopy (TEM, **Figure 3D**). The formation of nanoparticles allowed us to investigate the degradation of **P1** in aqueous buffer, to mimic physiological conditions. Since the disease microenvironment, such as the inflamed heart tissue, can be mildly acidic (pH 6.0~7.0),^40^ we examined nanoparticle degradation at three different pHs, including 6.0, 6.5 and 7.5 in buffer solution (**Figure 3E**, **Figure S8A**). At pH of 7.5, no significant change in ^31^P and ^1^H NMR spectra were observed after 10 days, suggesting the polymer backbone remained intact. In comparison, a minor peak at −0.45 ppm on ^31^P NMR was observed at more acidic pH, indicating partial backbone hydrolysis. Nevertheless, no signal from the final degradation product (i.e., phosphoric acid) was detected, indicating polymer was still the major species across the pH range. Despite this, formation of micron-size structures was observed via dynamic light scattering (DLS). These results indicate that the backbone integrity is crucial for maintaining a monodisperse micellar structure (**Figure S8B-C**).

To further understand this size change, another peptide brush polyphosphoramidate **P2** with a reduced P-N bond density (i.e., a hydrophobic block composed of MePTDO and NorPh random copolymer with same total DP) was prepared (**Figure S9A**). The incorporation of NorPh promoted the formation of smaller nanoparticles with a diameter of 20 nm (**Figure S9B**), possibly due to the higher T_g_ (120~150 °C depending on DP) associated with polynorbornene.^41^ Although the formation of phosphoric acid was still not detected (**Figure S9C-D**), the more intense ^31^P signal from the partially hydrolyzed intermediate suggests a faster backbone degradation. DLS analysis also revealed a polydisperse particle size distribution (**Figure S9E-F**). In particular, we observed two size transitions: one towards the formation of < 10 nm structures and the other towards the formation of microscale assemblies (> 10 μm). These findings provide insights into the particle degradation mechanism (**Figure S10**). We propose that the initial hydrolysis occurs at P-N bonds close to the hydrophilic peptide shell. After cleavage, the peptide brush is released, giving rise to a population with less than 10 nm diameters. The residual hydrophobic core then aggregates to generate the observed micron-sized assemblies. Therefore, under true physiological conditions, we would expect to see the complete disappearance of the microscale population as they degrade into water soluble small molecules.

### Enzyme Responsiveness, Cytocompability, and Hemocompatibility of Degradable Polyphosphoramidate Nanoparticles (DNPs)

An application of interest for these degradable polyphosphoramidate based nanoparticles (DNPs) is in targeted delivery. It is well established that MMPs are overexpressed in inflamed tissue, including the heart after acute MI. Previously, we have shown that nondegradable micellar nanoparticles assembled from MMP-responsive peptide polynorbornene amphiphiles can target and accumulate at the infarcted heart following systematic administration in a rat MI model.^1^ To examine whether these DNPs can be used as a targeted delivery platform, we first examined their enzyme responsiveness *in vitro*. A Cy5.5-labeled **P3** was prepared similarly and formulated into spherical micelles in buffer solution at 300 μM in respect of polymer (**Figure 4A-B**). This concentration was selected based on previously established procedures for *in vivo* intravenous (IV) injection.^1^ Upon overnight treatment with thermolysin, which is a robust thermostable alternative of MMP with which to test the system, a morphological switch into microscale assemblies was observed via TEM, indicating DNPs formulated from **P3** maintained the enzyme responsiveness as their polynorbornene based analog (**Figure 4C**).

**Figure 4.**
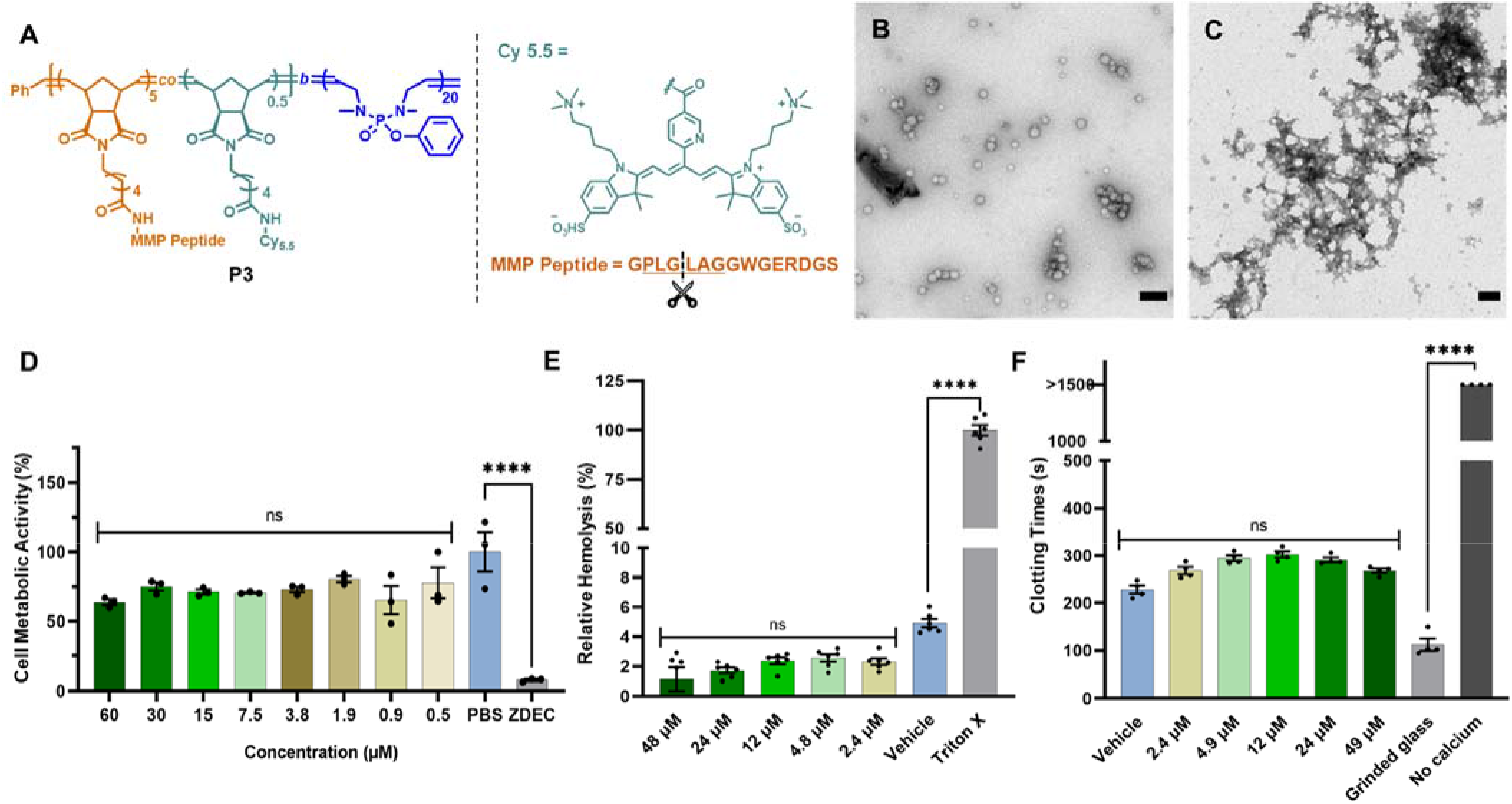
Enzyme responsiveness, cytocompatibility, and hemocompatibility of Cy5.5 labeled peptide-polyphosphoramidate nanoparticles (DNPs). (A) Chemical structure of peptide brush polyphosphoramidate **P3**. TEM images of (B) nanoparticles and (C) aggregates formation post incubation with thermolysin at 1:100 thermolysin:polymer for 24 h a 37 °C in DPBS. Scale bar: 100 nm. (D) Cytocompatibility of DNPs with L929 at various concentrations between 0.5 and 60 μM. For reference, the concentration of polymer in blood post IV injection is approximately 17.6 μM. (E) Percent hemolysis of red blood cells post incubation with DNPs at various concentrations. DPBS without Ca^2+^ and Mg^2+^ was used as the vehicle control, and 1% Triton X-100 as the positive control treatment. Absorbance was measured at 540 nm (n = 6). (F) Activated clotting times of whole human blood in the presence of DNPs at different concentrations. DPBS without Ca^2+^ and Mg^2+^ was used as the vehicle control, ground glass as the positive control, and no Ca^2+^ as the negative control treatment (n = 4). ns (p > 0.05), and **** (p ⩽ 0.0001) via Ordinary One-Way ANOVA. Values are displayed as mean ± SEM.

The cytocompatibility of these DNPs was then evaluated by incubating them at different concentrations with L929s, a line of murine fibroblasts used as a standard for cytotoxicity assays in accordance with ISO 10993 biocompatibility tests. The maximum concentration was chosen based off of the initial dilution of degradable NPs into the blood volume of a rat.^42^ Following treatment of L929s with various physiologically relevant concentrations of DNPs, we observed no significant decrease in metabolic activity compared to a healthy PBS control (**Figure 4D**).

To assess the hemocompatibility of DNPs, *in vitro* red blood cell (RBC) hemolysis and whole blood activating clotting time (ACT) assays were performed. In the RBC hemolysis assay, absorbance of isolated RBCs treated with DNPs was used to determine hemolytic activity. As compared to the vehicle (DPBS) control, the DNPs did not show any significant difference in percent hemolysis (**Figure 4E**, **Figure S11**). In the ACT assay, viscosity of human whole blood treated with DNPs over a range of dilutions was assessed by a Hemochron instrument. The DNPs performed similarly to the vehicle control (DPBS), though did show minimal anti-coagulative properties. However, the DNP treatments all had ACTs well below the range of the no calcium negative control (**Figure 4F**). These results suggest that the DNPs are hemocompatible and suitable for *in vivo* use.

### Targeted Delivery of DNPs in a Rat Myocardial Infarction Model

Following intravenous injection one-day post-MI, we observed strong DNP localization in the infarcted region of the heart. We detected regioselective accumulation in the infarcted region (**Figure 5A**) with very little retention in the border zone (**Figure 5B**) or remote myocardium (**Figure 5C**). Additionally, the morphology of the accumulation resembles a bolus, practically filling the injured myocardium with material. In the border zone and remote myocardium, DNPs appear more punctate and sparse.

**Figure 5.**
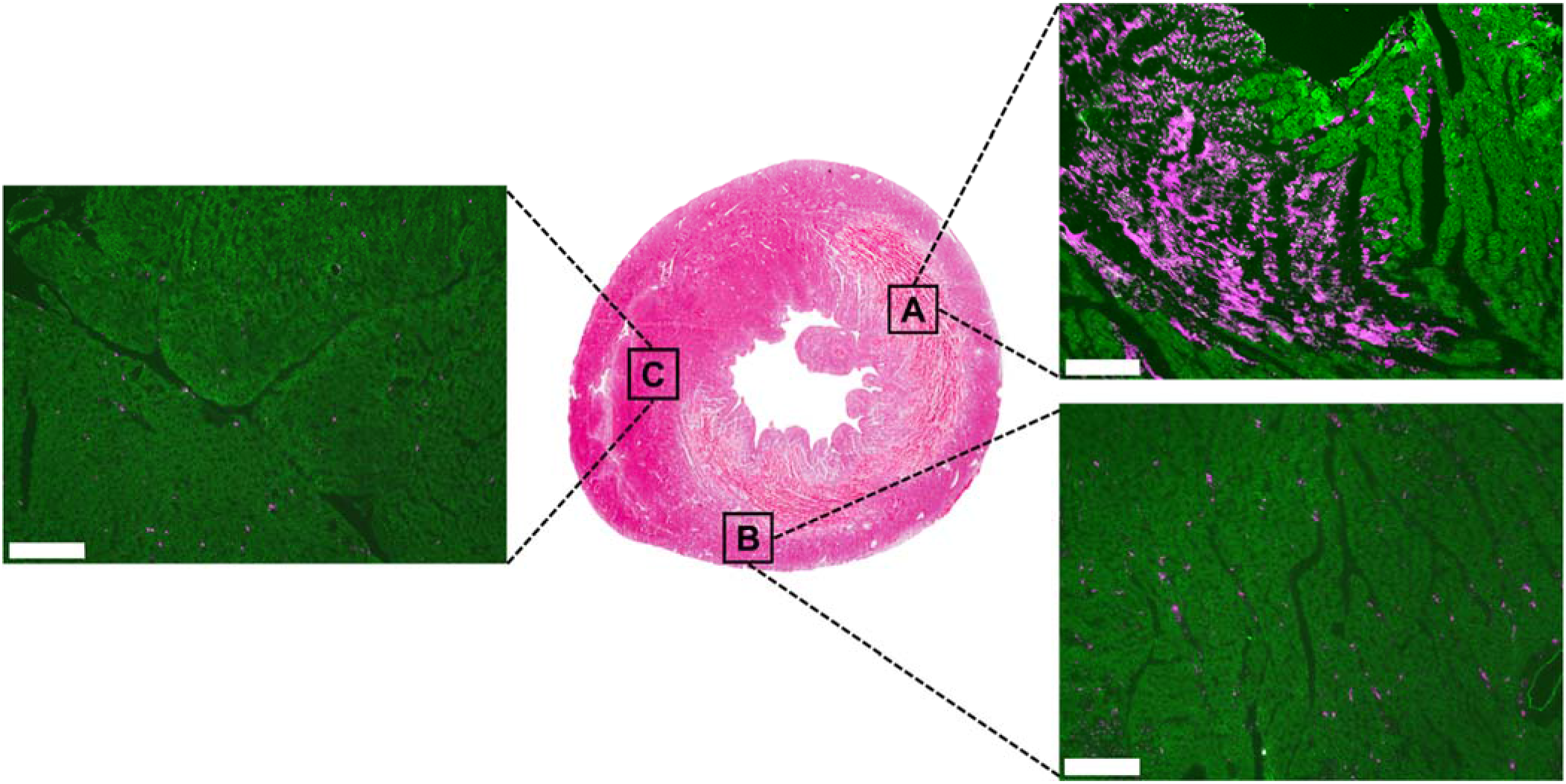
Selective nanoparticle accumulation in the infarcted heart. Middle panel shows the representative hematoxylin and eosin (H&E) stained heart section. (A-C) shows the corresponding fluorescence images of the neighboring section, which was stained for α-actinin (green), with Cy5.5-labeled DNPs displayed in red. (A) infarcted left ventricle, (B) borderzone, and (C) remote myocardium. Scale bar: 200 μm.

At one day post-injection, DNPs displayed strong accumulation in the heart, liver, and kidneys (**Figure 6A**). DNP intensity decreased over time out to 28 days-post injection (**Figure 6B-D**). Beyond targeting the infarcted myocardium, off-target accumulation in the liver and kidneys is typical for intravenously injected nanomaterials.^43^ Further, the DNP signal significantly decreased in the left ventricle (LV) at 7 days post-injection with further reduction in signal at 14 days (**Figure 6E**). This trend is also observed in the satellite organs; of particular interest is the decreased accumulation in the clearance organs, the liver and kidneys (**Figure 6F**).

**Figure 6.**
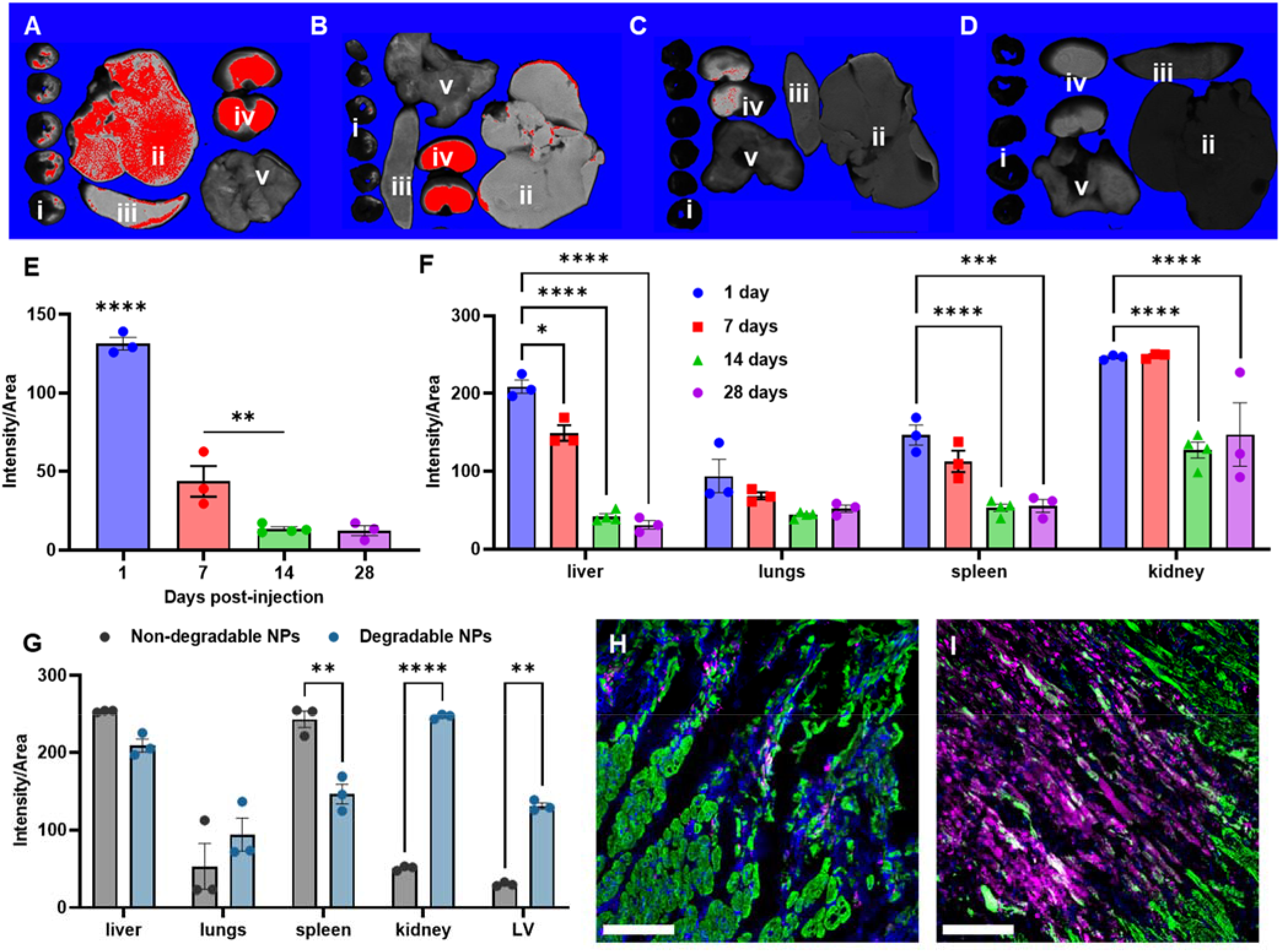
Degradable nanoparticle biodistribution over time. LiCor scans of the heart and satellite organs at (A) 1 day, (B) 7 days, (C) 14 days, and (D) 28 days post-injection of degradable nanoparticles (n = 3 at each timepoint, i: heart slices, ii: liver, iii: spleen, iv: kidneys, v: lungs). Quantification of nanoparticle signal in the (E) left ventricle and (F) satellite organs from LiCor organ scans. (G) Comparison of biodistribution of the original non-degradable platform to the degradable nanoparticles at 1 day post-injection. Immunohistochemistry for cardiomyocytes at 1 day post-injection for the (H) non-degradable and (I) degradable nanoparticle platforms. Heart sections were stained for nuclei (blue) and α-actinin (green), with DNPs displayed in red. Scale bar: 50μm. For Panel E: **(p ⩽ 0.01), and **** (p ⩽ 0.0001) via One-Way ANOVA with Tukey’s post-hoc test. For panels F and G: *(p ⩽ 0.05), **(p ⩽ 0.01), and **** (p ⩽ 0.0001) via Two-Way ANOVA with Šidák’s correction. Values are displayed as mean ± SEM.

Since P(MePTDO) completely hydrolyzed into small molecule fragments under acid catalyzed conditions, we hypothesized that DNPs can be gradually degraded and cleared from the body through the liver and kidneys. By 28 days there was trace DNP signal in the heart and satellite organs (**Figure 6E-F**), leading us to infer that the bulk of the initially injected material has been cleared from the body via bile and/or urine excretion.

When comparing the biodistribution of DNPs and their nondegradable counterparts (i.e., MMP responsive micellar nanoparticles assembled from peptide-polynorbornene amphiphiles) at 1 day postinjection, we observed significantly lower accumulation in the spleen accompanied by higher accumulation in the kidneys and LV (**Figure 6G**). Histology corroborated the differences in material accumulation in the infarcted LV. In addition, the analogous, non-degradable NPs accumulated in punctate aggregates (**Figure 6H**) whereas the DNPs appeared to accumulate to a much higher degree in the infarct (**Figure 6I**). Further imaging revealed colocalization of DNPs with cardiomyocytes in the necrotic infarct at 1 day post-injection (**Figure S12**), and with CD68^+^ macrophages at 14 and 28 days post-injection (**Figure S13**), suggesting macrophages are phagocytosing and clearing out the remaining DNPs.

## Conclusion

In summary, using PTDO as a starting point, we generated a library of diazaphosphepine-based cyclic olefins with different strain energies. Through this process, we identified a promising monomer, MePTDO, which can rapidly polymerize via ROMP at room temperature with > 84% monomer conversion in both DCM and DMF to afford well-defined polyphosphoramidates. These polymers underwent rapid degradation in acidic conditions with fast turnover from a partially hydrolyzed intermediate into phosphoric acid. We demonstrate that MePTDO is compatible with various norbornene monomers for copolymerizations, introducing degradability into otherwise nondegradable polymeric backbones. Most importantly for these studies, MePTDO can efficiently copolymerize with a norbornene functionalized with the peptide substrate of MMP, which enabled the fabrication of degradable peptide-polymer amphiphiles. The resulting copolymers assembled into biocompatible spherical micelles that responded to MMPs via phase transition and to acidic pH for degradation. As compared to their nondegradable counterparts, these degradable nanoparticles showed significantly increased accumulation in the infarcted heart at 24 hours post injection in a rat MI model. Moreover, instead of liver accumulation, these materials were predominantly detected in the kidney, suggesting renal clearance as the major route of clearance. At 14 days post injection, we observed a decrease in signal from the left ventricle of the heart, consistent with material degradation and clearance from the heart. Given the functional group tolerance of the ROMP initiators, a range of therapeutic compounds could be incorporated into future systems in the form of functionalized comonomers. These studies point a way forward to a generalizable and translational platform for drug delivery to treat different inflammatory diseases, where MMP upregulation is prevalent, especially in MI, where there remains an increasingly unmet clinical need for new treatments.

## Data Availability

All data supporting the findings of this study are available within the Article and its Supplementary Information.

## Acknowledgements

The authors are grateful for the support of the NIH through the NHLBI R01HL139001. Additionally, H.L.S. was supported by the Chemistry Biology Interfaces Training grant (5T32GM112584) and a National Institutes of Health Pre-Doctoral fellowship (1F31HL152610). Additional support was provided by the IMSERC at Northwestern University, which has received support from the Soft and Hybrid Nanotechnology Experimental (SHyNE) Resource (NSF ECCS-1542205), the State of Illinois, and the International Institute for Nanotechnology (IIN).

## Author Contributions

Y.L., H.L.S., K.L.C., and N.C.G. conceived this project. Y.L. conducted the monomer and polymer syntheses and performed the *in vitro* characterizations. H.L.S. conducted cytocompatibity and animal experiments. H.L.S. and K.W. performed *ex vivo* analysis. K.L.C. oversaw the *in vivo* study and C.L. conducted myocardial infarction surgeries. K.C. performed the hemocompatibity test and J.K. performed the transmission electron microscopy imaging. Y.L. and H.L.S. wrote the manuscript and received edits from K.L.C. and N.C.G. All the authors discussed the results and commented on the manuscript.

## Competing Interests

The authors declare no competing interests.

## Additional Information

Supplementary Information is available for this paper.

## Notes

### Competing Interest Statement

The authors have declared no competing interest.

