## Supporting Information for "Inflammation-Responsive Micellar Nanoparticles from Degradable Polyphosphoramidates for Targeted Delivery to Myocardial Infarction"

### 1. Materials

#### 1.1 Polymer and Nanoparticle Synthesis

Grubbs 2nd and 3rd generation catalysts were purchased from Sigma Aldrich. Amino acids used in Fmoc solid phase peptide synthesis were purchased from AAPPTec, ChemPep and NovaBiochem. Other reagents were purchased from commercial vendors and used as received unless otherwise noted. Phenyl, N-hydroxysuccinimide (NHS), and Cy5.5 labeled norbornene monomers were prepared as previously described.[1, 2] Anhydrous solvents, including dichloromethane (DCM) and dimethyformamide (DMF) were obtained from a Grubb's type solvent drying system prior to use. Flash column chromatography was performed using silica gel 60 (40-63 μm, 230-400 mesh, 60 Å) purchased from Fisher Scientific. Analytical thin-layer chromatography (TLC) was carried out on silica gel 60G F254 glass plates purchased from EMD Millipore and visualized by observation of fluorescence under ultraviolet light and staining with potassium permanganate ($KMnO_4$) as a developing agent. Dulbecco's phosphate buffered saline (without $Ca^{2+}$, $Mg^{2+}$) was purchased from Corning. Transmission electron microscopy (TEM) was performed on 400 mesh carbon grids purchased from Ted Pella, Inc.

#### 1.2 Enzyme-Responsiveness, Cytocompatibility and Hemocompatibility

Thermolysin was acquired from Promega, as a lyophilized powder. Murine L929s were obtained from Millipore. The alamarBlue™ reagent was ordered from Thermo. Citrated human whole blood was obtained from Innovative Research and stored at 1-6 °C prior to use.

#### 1.3 Animal Studies

Female Sprague-Dawley rats were purchased from Envigo and *ex vivo* organs were scanned on a LiCor Odyssey scanner. Antibodies were ordered from Signa (anti-$\alpha$-actinin) and BioRad (anti-CD68). Secondaries were obtained from Thermofisher (Alexa Fluor-488. Immunofluorescently stained slides were imaged using an upright Zeiss Fluorescent Microscope and a Keyence All-in-One Fluorescent Microscope.

#### 2. Instrumentation

**Nuclear Magnetic Resonance (NMR):** $^{1}H$-NMR and $^{31}P$-NMR spectra were recorded either on a 500 MHz Bruker Advance III HD system equipped with a TXO Prodigy probe or on a 400 MHz Bruker Advance III HD Nanobay system with SampleXpress autosampler. The residual solvent peaks were used as the reference signals (DMSO-$d_6$: δ 2.50 for $^{1}H$ NMR).

**Electrospray Ionization Mass Spectrometry (ESI-MS):** ESI-MS spectra were acquired on a Bruker AmaZon SL configured with an ESI source in both negative and positive ionization mode.

**Transmission Electron Microscope (TEM):** TEM samples were prepared by dropcasting 5 µL of sample on a TEM grid and wicking away excess liquid. TEM images were obtained using a JEOL 1230 transmission electron microscope operating at 120 keV equipped with a Gatan camera.

**Thermal Analysis:** DSC measurements were performed using TA DSC under nitrogen. Two thermal cycles (-50 to 200 °C) with heating and cooling rates of 10 °C/min were performed. Glass transition temperature was obtained from the second heating scan after the thermal history was removed.

**Size-Exclusion Chromatography (SEC):** SEC measurements were carried out in HPLC grade dimethylformamide (DMF) with 0.05 M LiBr at 60 °C on a Phenomenex Phenogel 5, 1K-75K, 300 x 7.80 mm column in series with a Phenomex Phenogel 5, 10K-1000K, 300 x 7.80 mm column. The detection system consisted of a L-2420 Hitachi UV-Vis Detector operating at 280 nm, a Wyatt Optilab T-rEX refractive index detector operating at 658 nm and a Wyatt DAWN® HELEOS® II light scattering detector operating at 659 nm. Absolute molecular weight and dispersity were calculated using the Wyatt ASTRA software with dn/dc values determined by assuming 100% mass recovery during SEC analysis.

**Analytical High-Performance Liquid Chromatography (HPLC):** Analytical HPLC analysis of peptides was performed on a Jupiter 4µ Proteo 90Å Phenomenex column (150 x 4.60 mm) using a Hitachi-Elite LaChrom L-2130 pump equipped with UV-Vis detector (Hitachi-Elite LaChrom L2420). The solvent system consists of (A) 0.1% TFA in water and (B) 0.1% TFA in acetonitrile.

**Preparative HPLC:** An Armen Glider CPC preparatory HPLC was used to purify peptides. The solvent system consists of (A) 0.1% TFA in water and (B) 0.1% TFA in acetonitrile.

**LiCor Odyssey:** Following nanoparticle injection and animal harvest, excised organs were scanned and quantified for fluorescent nanoparticle signal.

**Zeiss Fluorescent Microscope:** Fluorescently stained slides were imaged to determine material localization and degradation over time.

**Keyence BZ-X All-in-One Fluorescent Microscope:** Higher resolution imaging of fluorescently stained slides was performed to further investigate material retention and morphology in the heart.

**Zeiss LSM 780 Confocal Microscope:** Higher magnification images were taken to further determine the degree of localization and colocalization of nanoparticles with macrophages are later timepoints in the heart.

### 3. Experimental Procedures

#### 3.1 Monomer and Precursor Synthesis

**2-ethoxy-1,3,4,7-tetrahydro-1,3,2-diazaphosphepine 2-oxide (ETDO, 2)**

To 600 mL of dry DCM in ice/water bath, ethyl dichlorophosphate (1.27 g, 7.80 mmol, 1.0 equiv.) was added with stirring. 4-Dimethylaminopyridine (DMAP) (95.4 mg, 0.780 mmol, 0.1 equiv.) and triethylamine (8.20 mL, 58,6 mmol, 7.5 equiv.) were then slowly added. The solution was warmed up to r.t in 5 min and stirred for another 15 min until a slightly yellow color was observed. Cis-1,4-diamino-2-butene·2HCl (1.50 g, 9.37 mmol. 1.2 equiv.), which was prepared as previously described,[3] was separately dissolved in 50 mL of dry DCM and triethylamine (3.80 mL, 27.3 mmol, 3.5 equiv.). The mixture was slowly added to the ethyl dichlorophosphate solution over 30 min. The resulting mixture was stirred at r.t for 30 min and heated to reflux under $N_2$ for 24 h. The reaction was then cooled and concentrated under vacuum to about 100 mL. Water was added and the aqueous layer was extracted three times with DCM. The combined organic layer was dried over $MgSO_4$ and concentrated to dryness. The residue was purified via column chromatography (10% MeOH in DCM) to yield the product as a slightly yellow liquid (70 mg, 5%). $^1$H NMR (500 MHz, DMSO-*d6*) δ 5.53 (t, J = 2.2 Hz, 2H), 4.90 – 4.78 (m, 2H), 3.86 (dq, J = 8.1, 7.1 Hz, 2H), 3.42 – 3.35 (m, 4H), 1.20 (t, J = 7.0 Hz, 3H). $^{31}$P NMR (202 MHz, DMSO-*d6*) δ 20.58.

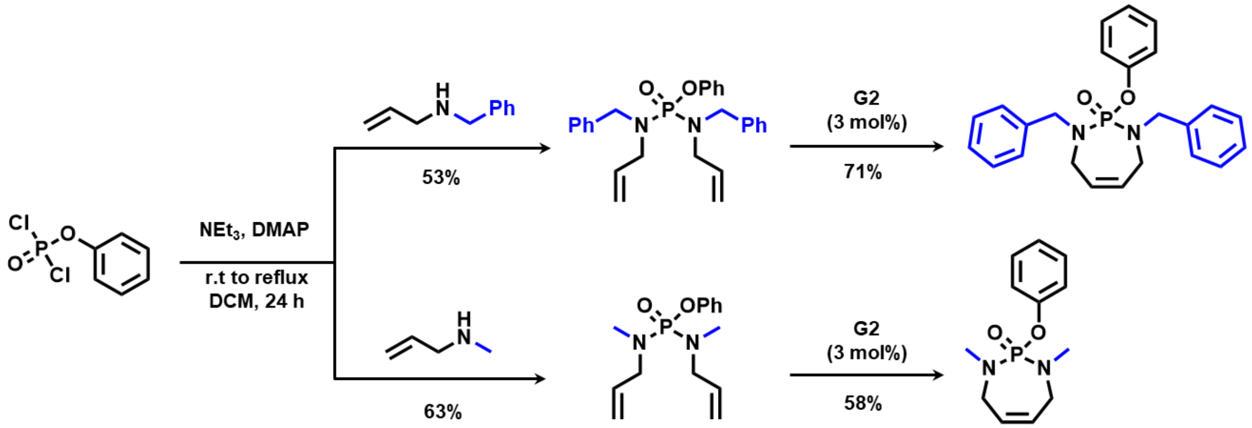

**Diene Precursor to 1,3-dimethyl-2-phenoxy-1,3,4,7-tetrahydro-1,3,2-diazaphosphepine 2-oxide (MePTDO, 3)**

To 18 mL DCM in ice/water bath, phenyl dichlorophosphate (3.1 g, 14.7 mmol, 1.0 equiv.), triethylamine (9.66 g, 95 mmol, 6.5 equiv.) and DMAP (180 mg, 1.4 mmol, 0.1 equiv.) were sequentially added. The reaction turned yellow and precipitation formation was observed. The reaction was warmed up to r.t and stirred for 15 min before dropwise N-allylmethylamine (2.15 g, 30 mmol, 2.05 equiv.) addition. The reaction mixture was brought to reflux and stirred overnight. The mixture was then partitioned between DCM and water. The aqueous layer was separated and washed with DCM. The organic layer was combined, washed with brine, and dried over $MgSO_4$. The crude was purified via column chromatography (1:1 EtOAc: Hex), affording product **2** as a colorless liquid. $^{1}$H NMR (400 MHz, DMSO-$d_6$) δ 7.42 – 7.31 (m, 2H), 7.23 – 7.11 (m, 3H), 5.70 (ddt, $J$ = 17.2, 10.1, 6.1 Hz, 2H), 5.26 – 5.08 (m, 4H), 3.71 – 3.45 (m, 4H), 2.58 (d, $J$ = 9.9 Hz, 6H). $^{31}$P NMR (162 MHz, DMSO-$d_6$) δ 14.53.

**1,3-dimethyl-2-phenoxy-1,3,4,7-tetrahydro-1,3,2-diazaphosphepine 2-oxide (MePTDO, 3)**

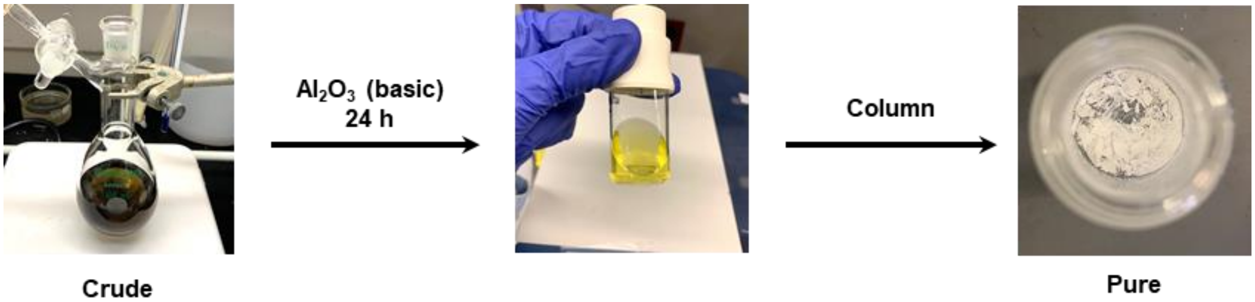

To 75 mL dry DCM in Schlenk flask, diene precursor **2** (540 mg, 1.93 mmol) was added. The solution was bubbled with $N_2$ for 10 min at r.t and heated to 40 ℃ for reflux. In a separate vial, **G2** (54.5 mg, 3 mol%) was added, degassed and then solubilized in DCM under $N_2$. The catalyst was then added to the diene solution and heated to reflux. After 5 min, a color change from purple to brown was observed. After 3 h, EVE was added to quench the catalyst and the reaction was allowed to cool to r.t in 30 min under air. A color change to green was observed, indicating catalyst deactivation. 50 wt. equiv. of basic aluminum oxide regarding to the crude was added to the reaction mixture and stirred overnight for catalyst absorptance. The aluminum oxide was removed through filtration and the residual solution was concentrated in vacuo. The crude was purified via column chromatography (3:1 EtOAc:Hex), affording the final product MePTDO as a white solid. $^{1}$H NMR (400 MHz, DMSO-$d_6$) δ 7.42 – 7.31 (m, 2H), 7.23 – 7.09 (m, 3H), 5.70 (t, $J$ = 2.6 Hz, 2H), 3.71 – 3.48 (m, 4H), 2.70 (d, $J$ = 9.3 Hz, 6H). $^{13}$C NMR (126 MHz, DMSO-$d_6$) δ 151.55, 151.50, 130.06, 127.83, 124.66, 120.74, 120.70, 46.89, 46.85, 36.26, 36.23. $^{31}$P NMR (162 MHz, DMSO-$d_6$) δ 17.40. ICP-MS analysis revealed a residual Ru of 0.54 ppm.

**Diene Precursor to 1,3-dibenzyl-2-phenoxy-1,3,4,7-tetrahydro-1,3,2-diazaphosphepine 2-oxide (BnPTDO, 4)**

The compound was synthesized using similar procedures as the diene precursor **2**. The crude was concentrated in vacuo and purified via column chromatography (4:1 Hex: EtOAc), giving the product as a colorless oil. $^{1}H$ NMR (500 MHz, DMSO-*d6*) δ 7.46 – 7.15 (m, 15H), 5.66 (ddt, J = 16.7, 10.2, 6.4 Hz, 2H), 5.20 – 5.06 (m, 4H), 4.30 – 4.11 (m, 4H), 3.60 – 3.41 (m, 4H). $^{31}P$ NMR (202 MHz, DMSO-*d6*) δ 13.54.

**1,3-dibenzyl-2-phenoxy-1,3,4,7-tetrahydro-1,3,2-diazaphosphepine 2-oxide (BnPTDO, 4)**

BnPTDO was synthesized following similar procedures for MePTDO. The **G2** catalyst was removed by overnight absorption with 50 wt. equiv. basic aluminum oxide. The aluminum oxide was removed through filtration and the residual solution was concentrated in vacuo before column chromatography (3:1 Hex:EtOAc) for purification. The product was isolated as a white solid. $^{1}H$ NMR (400 MHz, DMSO-$d_6$) δ 7.41 – 7.34 (m, 2H), 7.33 – 7.17 (m, 13H), 5.61 – 5.51 (m, 2H), 4.35 (dd, $J$ = 15.4, 10.0 Hz, 2H), 4.17 (dd, $J$ = 15.4, 6.9 Hz, 2H), 3.72 – 3.43 (m, 4H). $^{31}P$ NMR (162 MHz, DMSO-$d_6$) δ 17.15.

**3.2 ICP-MS for the Measurement of Residual Ru**

200 µL HNO3 and 200 µL HCl was added to 17.05 mg MePTDO. The mixture was heated in 65 °C water bath for 3 hours for acid digeastion. The resulting solution was diluted to 10 mL with millipore water and ran versus 5 ppb max std curve on ICP-MS to detect the residual Ru. The Ru level in MePTDO was determined to be 538.992 ppb (0.54  ppm).

**3.3 Synthesis of Peptide Substrate of Matrix Metalloproteinase (MMP) and Peptide-Conjugated Norbornene Monomer (NorMMP)**

Peptides with the amino acid sequence, GPLGLAGGWGERDGS, were synthesized on rink amide 4-methyl benzylhydrylamine (MBHA) resin via Fmoc-based solid phase peptide synthesis. The underlined amino acids represent an MMP-2 and -9 recognition sequence. The resin was allowed to swell in DMF for 2 h. Fmoc deprotection was performed by agitating resin in 20% 4-

methylpiperidine in DMF for 5 draining, and repeating this procedure for another 15 min. Amino acid couplings were carried out for 45 min per amino acid using N,N,N',N'-tetramethyl-O-(1H-benzotriazol-1-yl)uronium hexafluorophosphate (HBTU) and N,N-diisopropylethylamine (DIPEA) (resin/amino acid/HBTU/DIPEA 1:3:3:6). Final peptides were cleaved from resin by treatment with trifluoracetic acid (TFA), triisopropyl silane (TIPS), dithiothreitol (DTT), and water (TFA/TIPS/DTT/$H_2O$ 88% v/v:2% v/v:5% w/v: 5% w/w) for 2 hours. Peptides were then precipitated in cold diethyl ether and centrifuged at 10000 rpm for 10 min. This procedure was repeated twice. The precipitated peptide was dried in vacuo to give the crude product. The purity of the crude was examined via reverse HPLC running under a mixture of Buffer A (0.1% TFA in $H_2O$) and Buffer B (0.1% TFA in ACN). The peptide afforded a signal at 14 min elution time over a gradient of 15~40% B in 30 min (detection at 214 nm). Same gradient was adopted for peptide purification by a preparation grade HPLC. The purified peptide was dried via lyophilization, giving a white solid. ESI-MS: calculated for $C_{61}H_{94}N_{20}O_{20}$ $[M+H]^+$ 1427.7; found $[M+H]^+$ 1427.7 and $[M+2H]^{2+}$ 714.4.

With the MMP peptide still on resin, NorAHA (3 equiv.), HBTU (3.0 equiv.) and DIPEA (6.0 equiv.) in DMF were added. After 24 h, the reaction solution was drained, and the resin was rinsed with DCM three times. Peptide monomer was cleaved by treating with 88%: 5%: 5%: 2% TFA: TIPS: DTT: $H_2O$ for 3 hours. The resulting solution was concentrated in vacuo before addition of cold diethyl ether for peptide precipitation. The crude product was dried and purified via a preparation grade HPLC over a gradient of 20~60% Buffer B in 30 min. The product eluted out at 16.5 min (~ 40% B) was collected and dried via lyophilization to afford the pure NorMMP monomer as a white powder. ESI-MS: calculated for $C_{76}H_{111}N_{21}O_{23}$ $[M+H]^+$ 1687.82; Found 1688.09.

#### 3.4 Polymer Synthesis

##### Synthesis of P(MePTDO) Homopolymer

Dry DCM was freeze-pump-thawed three times to remove air before moving into the glovebox under $N_2$. **G3**, $(IMesH_2)(C_5H_4NBr)_2(Cl)_2Ru{=}CHPh$, and MePTDO were weighed into separate vials and moved into the glovebox. In the box, **G3** and MePTDO were dissolved with DCM and then mixed for polymerization. An immediate color change from green to brown was observed, indicating the initiation of polymerization. The sealed reaction vial was then moved out of the box.

After designated reaction time, ethyl vinyl ether (EVE) was added and stirred for 10 min to quench the catalyst. Polymer was collected as the solid post precipitation into cold diethyl ether and dried in vacuo. $^{1}$H NMR (400 MHz, DMSO-$d_6$) δ 7.40 – 7.26 (m, 2H), 7.13 (dd, $J$ = 15.6, 7.8 Hz, 3H), 5.46 (t, $J$ = 3.4 Hz, 2H), 3.68 – 3.43 (m, 4H), 2.52 (s, 6H). $^{13}$C NMR (126 MHz, DMSO-$d_6$) δ 151.49, 151.44, 130.00, 129.85, 129.83, 124.58, 120.68, 120.64, 50.46, 50.43, 33.37, 33.34. $^{31}$P NMR (162 MHz, DMSO-$d_6$) δ 14.54 (d, $J$ = 16.1 Hz).

**Synthesis of Norbornene and MePTDO Block Copolymer:**

**G3**, norbornene and MePTDO were weighed into separate vials and moved into the glovebox under $N_2$. The reagents were dissolved in dry and air-free DCM. The norbornene solution was added to **G3** for polymerization. After 45 min, the MePTDO solution was added to the reaction mixture. The polymerization was quenched with EVE post 1.5 h and the reaction crude was precipitated into cold diethyl ether for monomer removal. The copolymer was collected as the solid. Note: To synthesize the peptide functionalized polyphosphoramidates, polymerizations were performed in DMF with 1M LiBr to aid peptide solubilization.

**Synthesis of Norbornene and MePTDO Random Copolymer**

The random copolymer was synthesized following similar procedures as the block copolymer. Norbornene and MePTDO were weighed and solubilized in the same vial before adding to the **G3** solution. The polymerization was quenched with EVE after 1.5 h of reaction and precipitated into cold diethyl ether to collect the polymer.

**3.5 Acid Catalyzed Monomer and Polymer Degradation**

**Accelerated Degradation for NMR Analysis**

MePTDO or P(MePTDO) was solubilized in DMSO-*d6* at 4 mg/mL. 4 M DCl in $D_2O$ was added, giving a 0.25 M DCl solution. The solution was transferred to NMR tube and the degradation was monitored on a daily base.

**Acclerated Degradation of Polymer for SEC-MALS Analysis**

P(MePTDO) was solubilized in DMF at 4 mg/mL. 4 M HCl was added, giving a 0.25 M HCl. An aliqot of the solution was removed every 24 h, dried with $MgSO_4$ and centrifuged. The top clear layer was used for SEC-MALS analysis. For copolymer, 0.5 M HCl in DMF was used to accelerate the degradation.

##### 3.6 Nanoparticle Formulation and Degradation

P(NorMMP)-*b*-P(MePTDO) (with or without the NorCy5.5, with or without NorPh as comonomer) was solubilized in DMSO at 3 mg/mL. Millipore water was added to the polymer solution with stirring using synringe pump at 0.3 mL/h until reaching 30% v/v water/DMSO. After complete water addition, the mixture was transferred into SnakeSkin™ Dialysis Tubing (7K MWCO) and dialyzed against water for 48 h with three water changes. The solution was then dialyzed into 1X Dulbecco's phosphate-buffered saline (DPBS) without $Ca^{2+}$ and $Mg^{2+}$, filtered through 0.22 µm PES membrane and concentrated to afford a nanoparticle solution with 300 µM polymer. Note: the concentration is calculated based on 1) initial polymer weight to final volume; or 2) Cy5.5 UV absorbance.

Peptide functionalized polyphosphoramidate was first assembled into nanoparticles in 1X DPBS (pH = 7.5), giving a final concentration of 4 mg/mL. An aliquot was removed, lyophilized dry and redissovled in DMSO-*d6* for NMR measurement (t = 0 point). The remaining solution was divided into three portions. The solution pHs were ajusted with 1 M HCl and confirmed by a pH meter to obtain the nanoparticle samples at pH = 6.0, 6.5 and 7.5. The particle morphology was monitored daily via DLS. At day 10, the samples were lyophilized dry and redissolved in DMSO-*d6* for NMR measurement.

##### 3.7 Enzyme Responsiveness

Thermolysin stock solution (1 mg/mL, 29 µM) was prepared in 1X DPBS added to Cy5.5 labeled P(NorMMP)-*b*-P(MePTDO) nanoparticles, giving 1 µM: 100 µM thermolysin: NPs in DPBS. The mixture was incubated at 37 °C for 24 h before cooling down to 2 °C to deactivate the enzyme. The nanoparticles before and after thermolysin treatment were imaged with TEM.

##### 3.8 Ring Strain via Density Functional Theory Calculation

Density functional theory (DFT) calculations were performed using Spartan 14.[4, 5] Geometry optimizations of reactants and product were performed at the B3LYP/6-31 g(d) level of theory. The conventional B3LYP functional has been proven to generate adequate geometries but perform poorly in energy calculations. Therefore, the energies were refined with the ω B97X-D functional including dispersion corrections and a 6-31 g(d) basis set. The enthalpy or heat of formation has been estimated as the energy difference between the total energy of the ring-opened molecule and the energy of the isolated reactants (PTDO + ethylene). DCM was used as the solvent for geometry optimizations and energy calculations.

##### 3.9 Cytocompatibility

For cytocompatibility assessment, murine fibroblast cells (L929) were used in accordance with the UNI ISO 10993/2009 for cytotoxicity assays. Cells were plated and left to adhere overnight. Following cell adhesion, degradable nanoparticles (DNPs) were added at physiologically relevant concentrations spanning 60-0.5μM with PBS and zinc diethyldithiocarbamate (ZDEC) serving as positive and negative controls, respectively. Treated cells were then incubated for 24 hours before performing an Alamar Blue assay to evaluate their metabolic activity. All treatments were normalized to the healthy PBS control.

##### 3.10 Hemocompatibility

Hemocompatibility analysis was performed similarly as described by Carlini et al.[6, 7]

###### Nanoparticle Dilutions for Hemocompatibility

According to previously optimized surgical procedures, 1 mL nanoparticle solution at 300 μM regarding polymer can be injected intravenously into rat with myocardial infarction. Under the assumption that a 250 g rat has 16 mL of blood, polymer concentration in bloodstream is approximately 17.6 μM. Therefore, degradable nanoparticles (DPNs) stock solutions at 600, 300, 150, 60 and 30 μM in 1X DPBS without $Ca^{2+}$ and $Mg^{2+}$ were prepared using serial dilutions.

###### Hemolysis of Red Blood Cells (RBCs)

Stocks of DNPs were added to the isolated red blood cells (RBC) so that final polymer concentrations were 48, 24, 12, 4.8, 2.4 μM. 1X DPBS without $Ca^{2+}$ and $Mg^{2+}$ was used as the vehicle control and 1% Triton X-100 as the positive control treatment. RBCs were isolated from 40 mL of citrated human whole blood via centrifugation at 500 x g for 5 min. Supernatant was removed and replaced with 150 mM NaCl solution. The RBCs were gently mixed and re-isolated with centrifugation. Gentle washes with DPBS were repeated three times. Once isolated, the RBCs were diluted 1:50 in 1X DPBS and gently mixed. RBCs (184 μL) were treated with DNP dilutions (16 μL) in a clear 96-well plate and incubated for 1-hour at 37 °C (n=6). Due to the innate color of the dye-labeled NPs, 16 μL of DNP dilutions was incubated with 184 μL of DPBS in parallel to adjust absorbance values (n=3). Plates were centrifuged at 500 x g for 10 min to form a pellet of intact RBCs. 100 μL of the supernatant from each well was carefully transferred to a fresh, clear 96-well plate. Absorbance measurements were taken to determine hemolysis and recorded at 405 nm, 450 nm, and 540 nm as well as representative full spectra from each experimental group. Plate reader measurements were conducted on an EnSpire Multimode Plate Reader. Hemolysis was determined by correcting for the background absorbance of the plate and respective absorbance from each sample group of parallel DNPs in DPBS and then normalized to 1% Triton X-100-treated RBCs (representing 100% hemolysis). Statistical comparisons were performed using a one-way ANOVA.

**Activated Clotting Time (ACT) Assay with Whole Human Blood**

Stocks of DNPs were added to whole human blood so that final polymer concentrations were 49.0, 24.5, 12.2, 4.91, 2.45 μM. 1X DPBS without $Ca^{2+}$ and $Mg^{2+}$ was used as the vehicle control, ground glass as the positive control, and no $Ca^{2+}$ as the negative control treatment. Using a calibrated Hemochron 801 instrument, activated clotting time of the DNPs in recalified citrated human whole blood was assessed. Briefly, Hemochron P214 tubes with glass beads were warmed and 4 μL of $CaCl_2$ (2.2 M) and 36 μL of DNP stock or control additive was added to each tube, gently mixed, and allowed to incubate for 30 s at 37 °C (n=4). Citrated human whole blood (400 μL) was added to each tube (t=0), mixed by hand for 10-15 seconds, and inserted into the instrument. Clot formation was determined by the displacement of the magnet within the tube. The time to clot formation was recorded for each sample. The negative controls (blood samples without calcium) exceeded the maximum time range of the instrument (>1500 s). The ACTs for the DNPs are reported in comparison to the positive and negative controls.

##### 3.11 Animal Studies

###### Surgical procedures and IV injection

All procedures in this study were approved by the Committee on Animal Research at the University of California, San Diego and the Association for the Assessment and Accreditation of Laboratory Animal Care. Female, Sprague Dawley rats (225 – 250g) underwent ischemia-reperfusion (IR) procedures via left thoracotomy and temporary occlusion of the left anterior descending artery for 35 minutes[8]. One day post-MI, animals were anesthetized using isoflurane and randomly intravenously injected with 1 mL of NPs (300µM) and harvested at 1, 7, 21, and 28 days post-injection (n = 3 for each timepoint). Animals were euthanized via overdose of pentobarbital (200 mg/kg) and the satellite organs (kidney, spleen, lungs, and liver) were collected for LiCor analysis. The heart was excised and matrix sliced into 5 sections for LiCor analysis and immunohistochemistry.

###### LiCor Odyssey whole organ scanning and quantification

Organs were kept on ice until scanning. A clear transparency was placed on the scanner and the organs were arranged and scanned at intensity level 1 at the 700-nanometer wavelength with an offset of 1 millimeter. Scanned images were analyzed using a custom MATLAB script where individual organs were outlined and analyzed for Cy5.5 fluorescence intensity per area.

###### Immunohistochemistry (α-actinin, CD68)

Following euthanasia, hearts were sliced into 5 sections and embedded in OCT for cryosectioning. Hearts were sectioned to a thickness of 10 µm sections and stained with anti-α-actinin (1:75 dilution, Sigma) and Alexa Fluor-488 (1:500 dilution, ThermoFisher). To identify macrophages, sections were stained with anti-CD68 (1:100 dilution, BioRad) and Alexa Fluor-488 (1:500 dilution, ThermoFisher). Slides were imaged using a Keyence BZ-X Series all-in-one fluorescent microscope and a Zeiss LSM 780 confocal microscope.

#### 3. Supporting Table and Figures

**Table S1.** Experimental and DFT calculated ring strain energy.

| Name | DFT Ring Strain (kcal/mol) | Experimental Ring Strain (kcal/mol) |
|---|---|---|
| **ETDO** | 3.44 | --- |
| **MePTDO** | 7.21 | --- |
| **BnPTDO** | - 0.50 | --- |
| **Norbornene** | 16.2 | 27.2[9] |

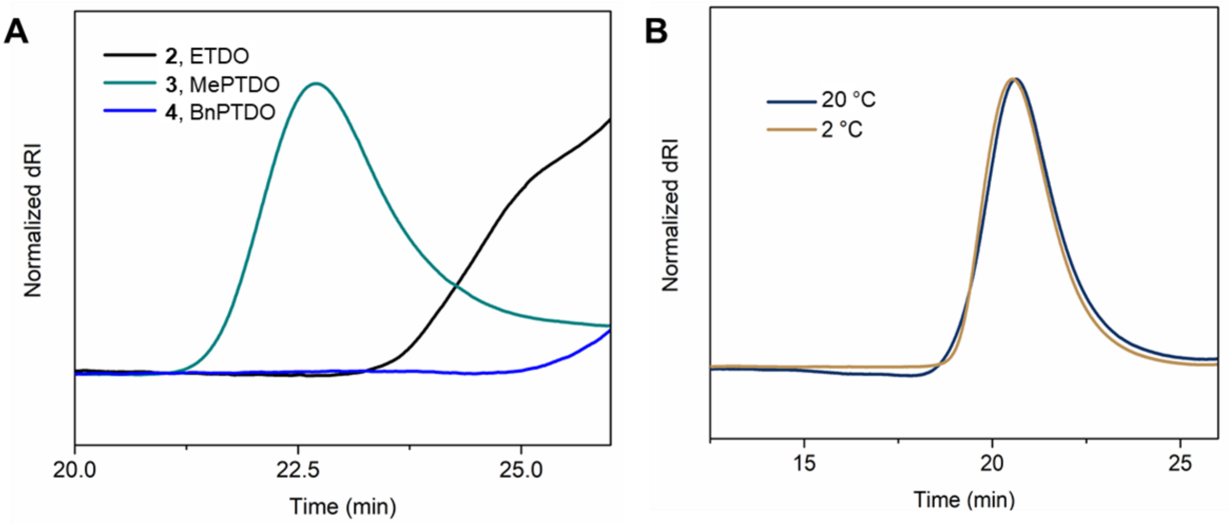

**Figure S1.** (A) SEC traces of test polymerizations of monomer **2** (EtDO), **3** (MePTDO), and **4** (BnPTDO). The reactions were performed using the previously reported condition: 2 °C, $[M]_0$ = 0.5 M, 5 h in 10/90 v/v MeOH/DCM. (B) SEC traces of **3** (MePTDO) polymerizations at 2 °C and room temperature (20 °C).

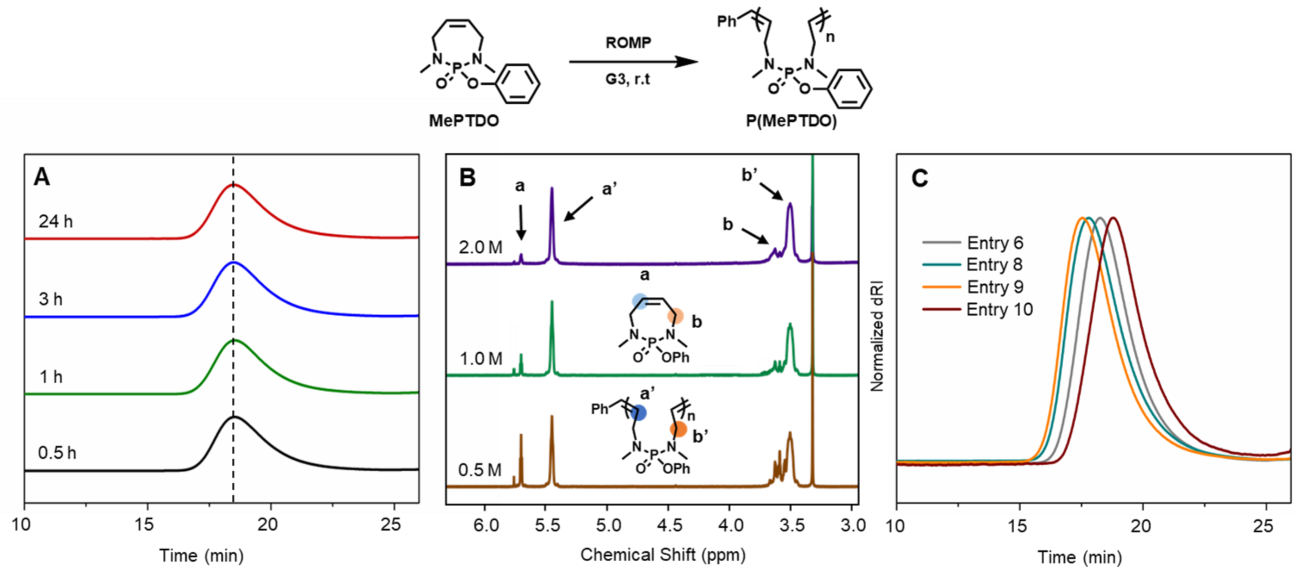

**Figure S2.** Condition screening for MePTDO homopolymerizations. (A) SEC traces of polymerizations terminated at different times. (B) Crude $^{1}$H NMR spectra of the polymerizations with different initial monomer concentrations $[M]_0$. Alkene signals from monomer and polymer were labeled to demonstrate the improved monomer conversion with higher $[M]_0$. (C) SEC traces of polymerizations with different $[M_0]$: **G3** ratios in accordance with **Table 1**.

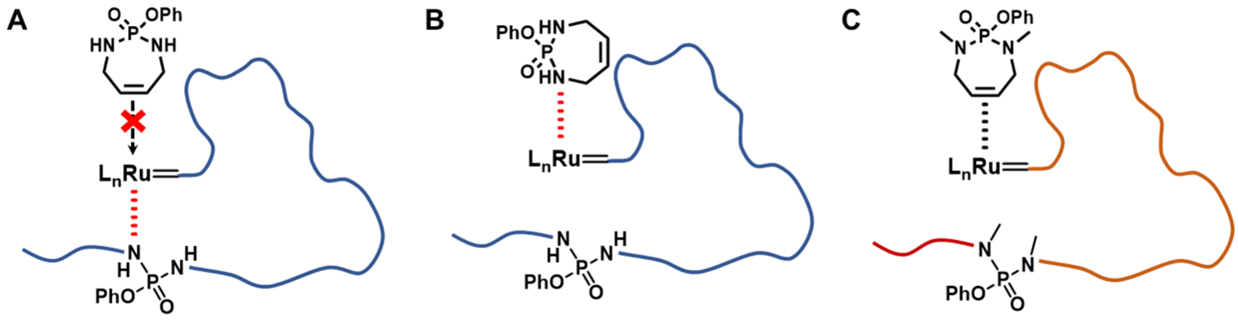

**Figure S3.** Proposed mechanism for competitions between amine and olefin on incoming monomer for the coordination site of propagating Ru carbene. For PTDO: (A) The secondary amine on the propagating chain rewinds and coordinates to the Ru carbene, blocking the incoming monomer. (B) The secondary amine on PTDO coordinates onto the Ru carbene. For MePTDO: (C) Methylation on amine reduces the unwanted amine-catalyst interaction, allowing proper monomer coordination onto the Ru carbene for subsequent ring opening.

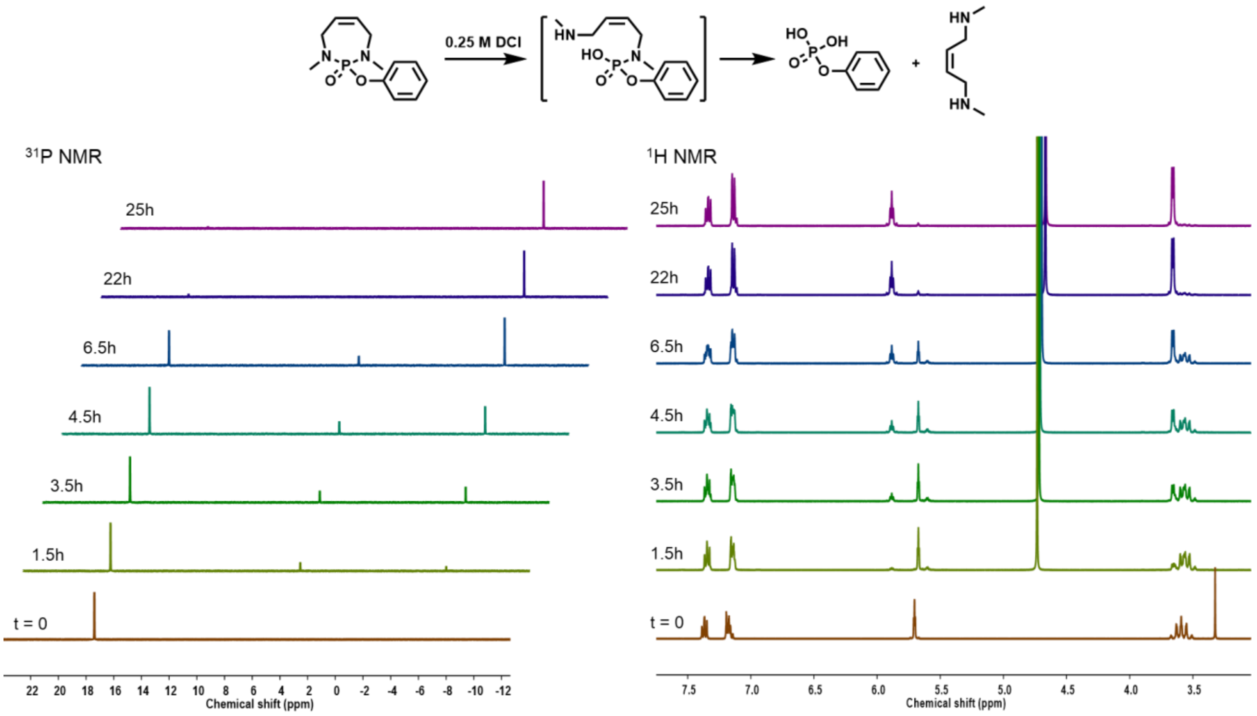

**Figure S4.** $^{31}P$ and $^{1}H$ NMR spectra of MePTDO degradation over time.

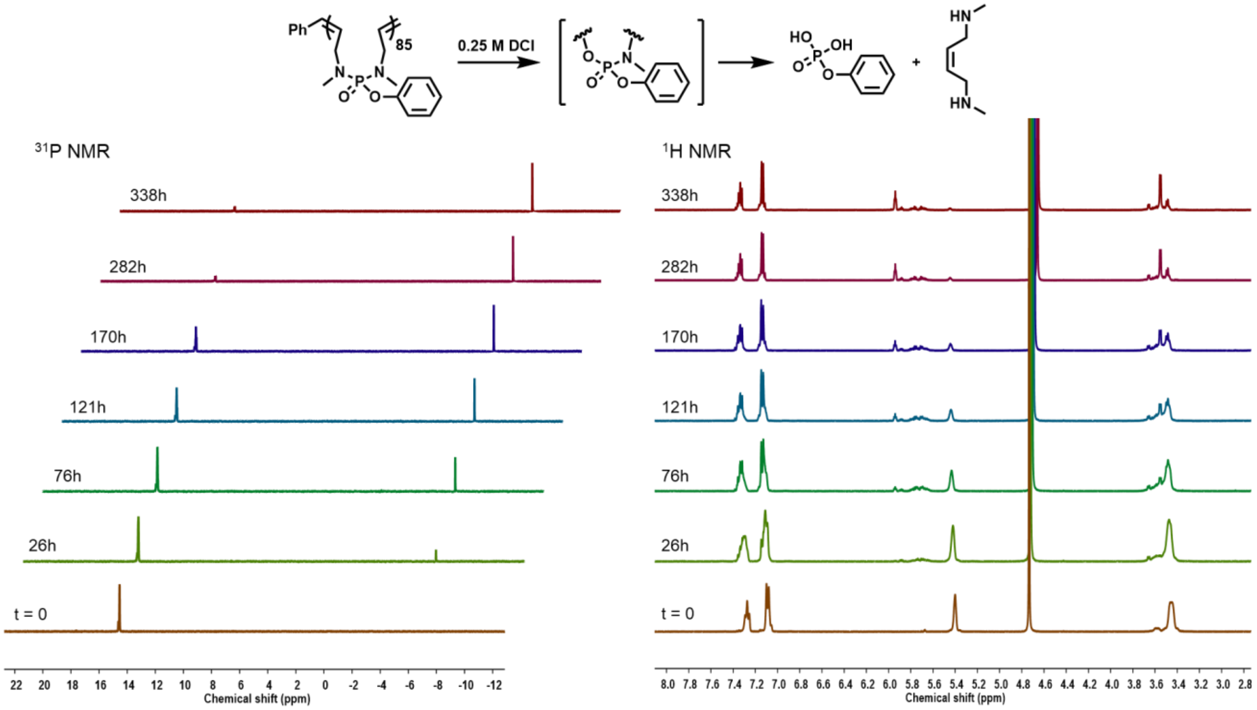

**Figure S5.** $^{31}P$ and $^{1}H$ NMR spectra of P(MePTDO) degradation over time.

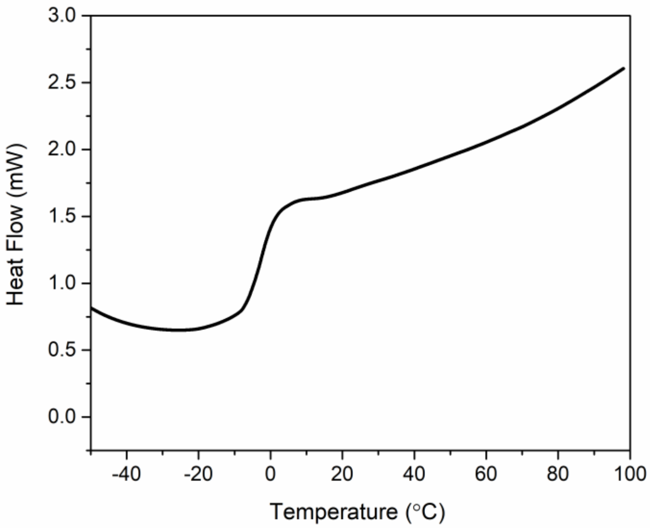

**Figure S6.** DSC measurement of P(MePTDO) (DP = 88). The graph depicts endothermic heat flow. A glass transition at -3 °C was detected.

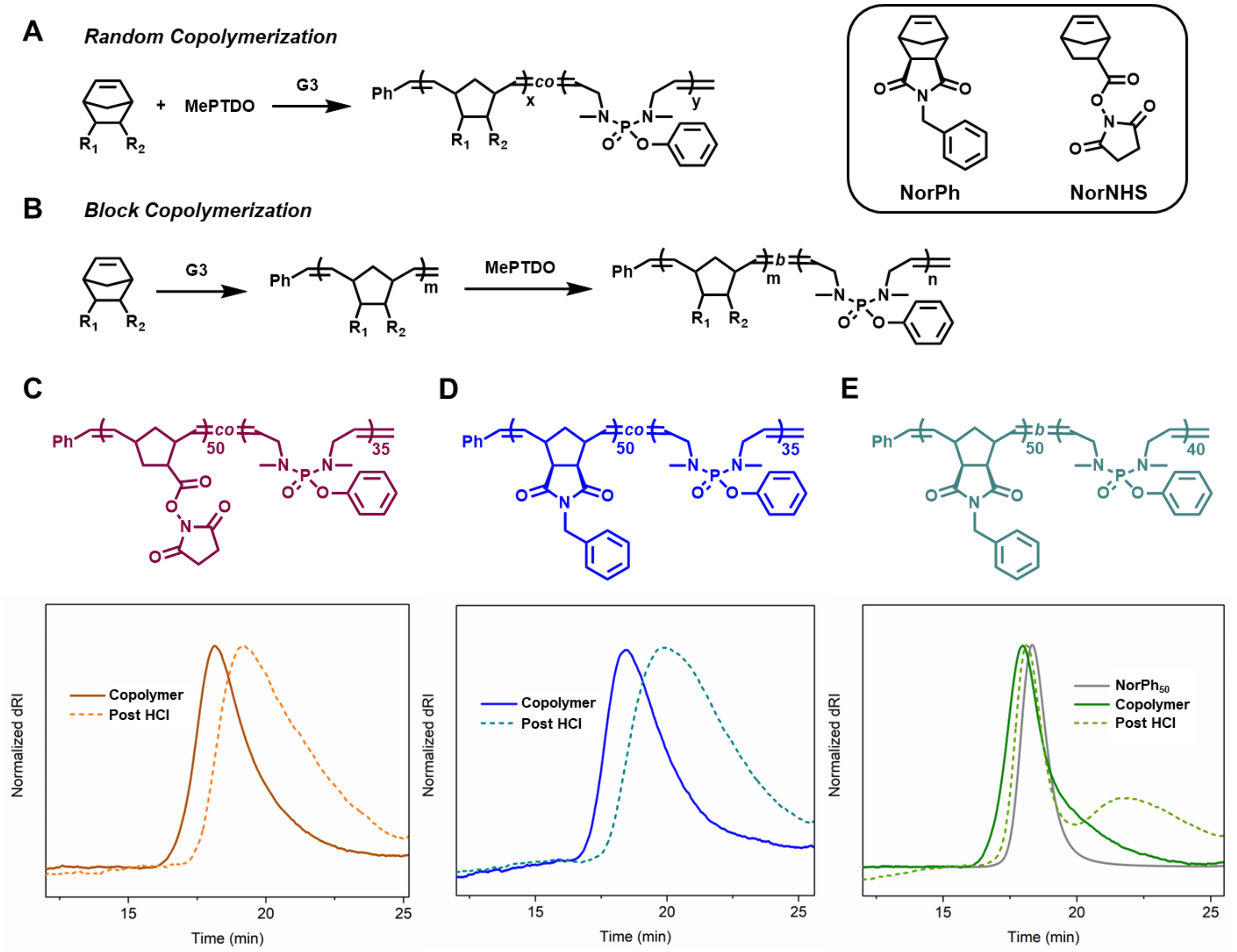

**Figure S7.** MePTDO copolymerization with NorPh and NorNHS. The reactions were performed in DCM at 20 °C with 50:50:1 MePTDO: norbornene: **G3** and $[MePTDO]_0$ of 0.5 M. Synthetic scheme for (A) random copolymerization and (B) block copolymerization. SEC traces of the random copolymers (C) $NorNHS_{50}$-*co*-$MePTDO_{15}$ and (D) $NorPh_{50}$-*co*-$MePTDO_{35}$ before and after acid treatment. (E) SEC traces of the block copolymer $NorPh_{50}$-*b*-$MePTDO_{40}$ before and after acid treatment. A polynorbornene control ($NorPh_{50}$) was included as the reference. Degradation condition: 0.5 M HCl in DMF for 48 h.

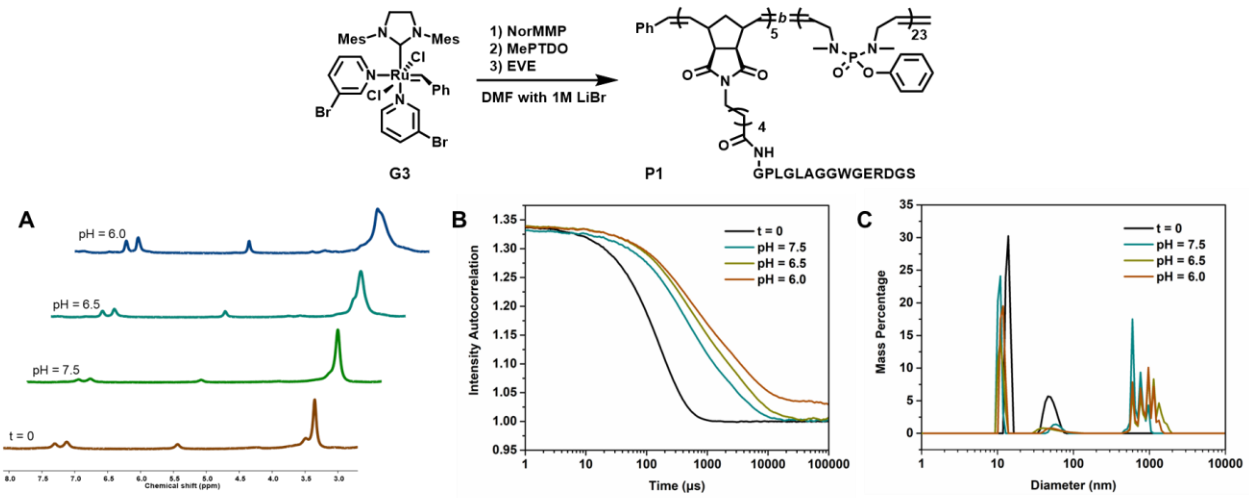

**Figure S8.** Synthesis and degradation of **P1** and formulated nanoparticles. (A) $^1$H NMR spectra of **P1** after 10 days in DPBS buffer at pH = 6.0, 6.5 and 7.5. Spectrum at t = 0 was included as the reference. (B) Autocorrelation curves and (C) DLS analysis of **P1** nanoparticles before and after 15 days in buffer solutions at different pHs.

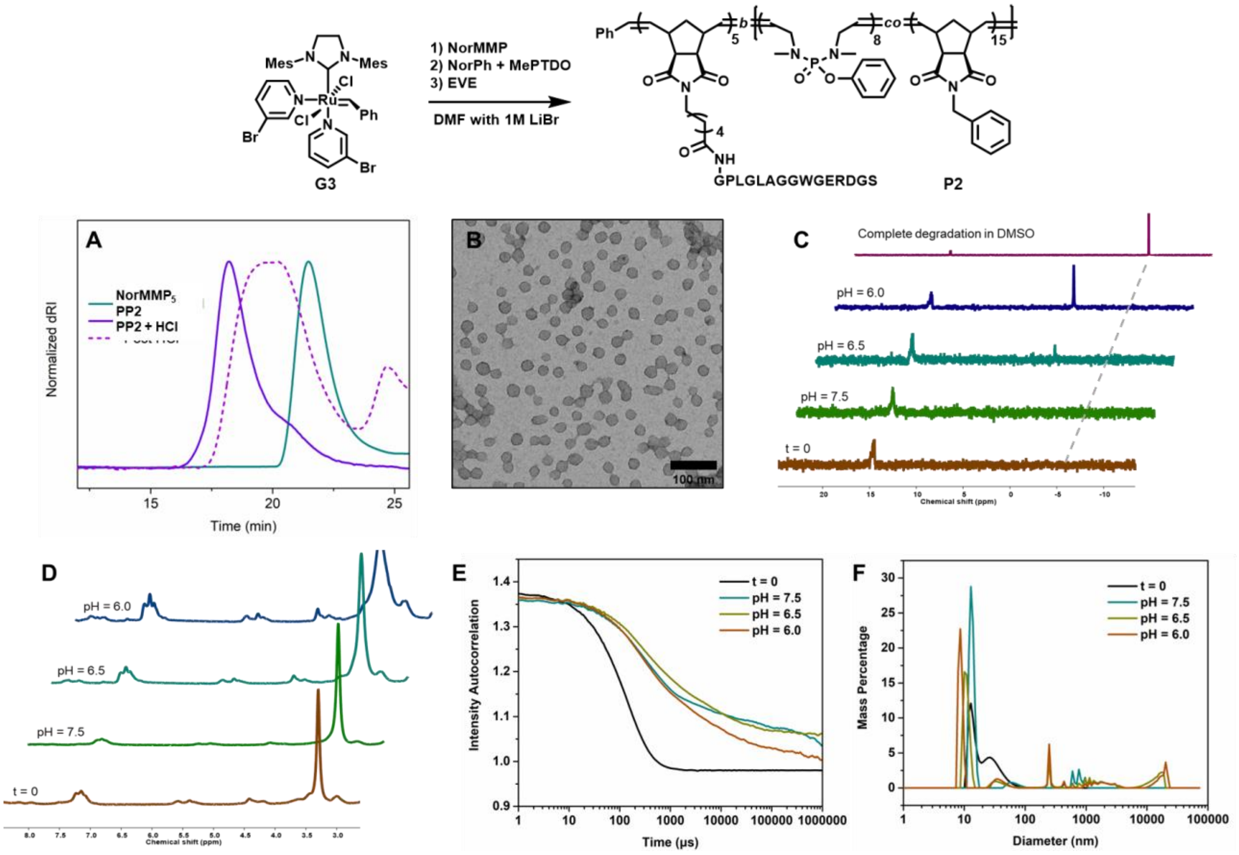

**Figure S9.** Synthesis and degradation of **P2** and nanoparticles. (A) SEC traces of **P2** before and after accelerated degradation in 0.5 M HCl in DMF. $NorMMP_5$ was used as the control. (B) TEM image of **P2** assembly into micelles. (C) $^{31}P$ and (D) $^{1}H$ NMR spectra of **P2** post 10 days in DPBS buffer at pH = 6.0, 6.5 and 7.5. Spectra at t = 0 and post accelerated degradation were used as reference. (E) Autocorrelation curves and (F) DLS analysis of **P2** particles before and after 15 days in buffer at different pHs.

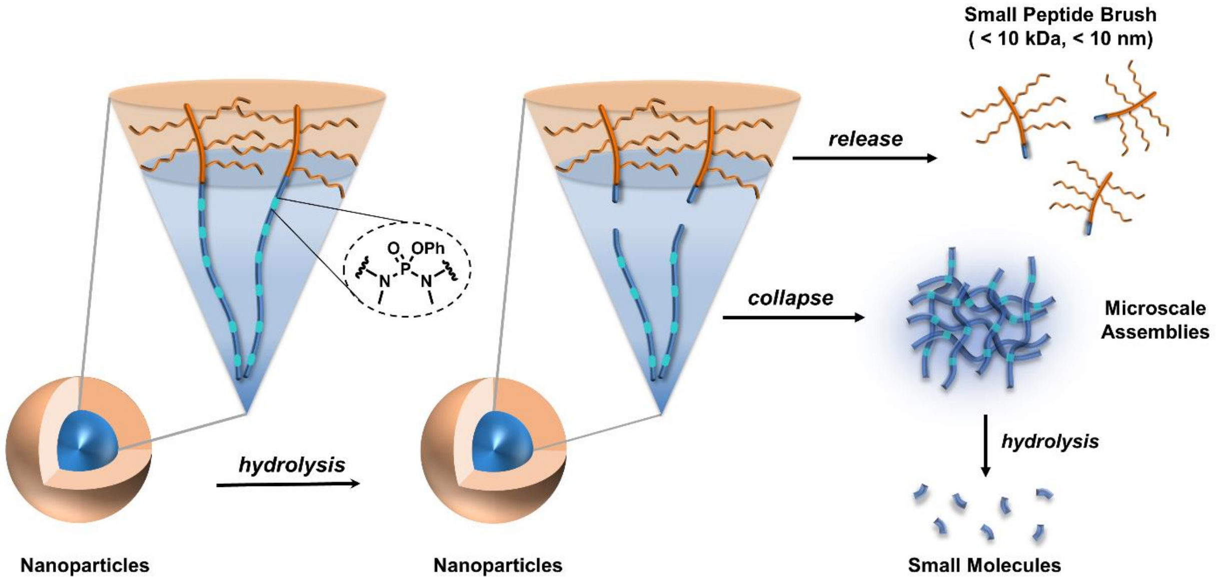

**Figure S10.** Proposed mechanism of **P3** nanoparticles degradation in aqueous environment.

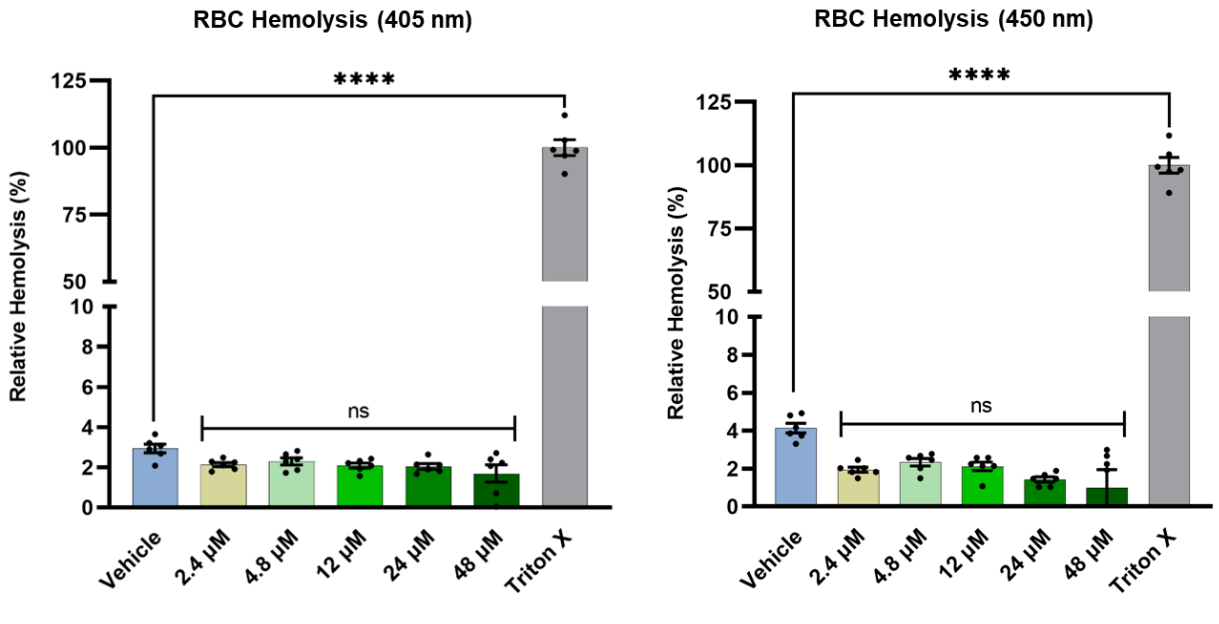

**Figure S11.** Red blood cell hemolysis post incubation with nanoparticles assembled from **P3** at various concentrations. (n = 6 per group). The absorbance was detected at 405 nm (left) and 250 nm (right). ns ($p > 0.05$), and ****$p \leqslant 0.0001$ via ordinary one-way ANOVA. Values are displayed as mean ± SEM.

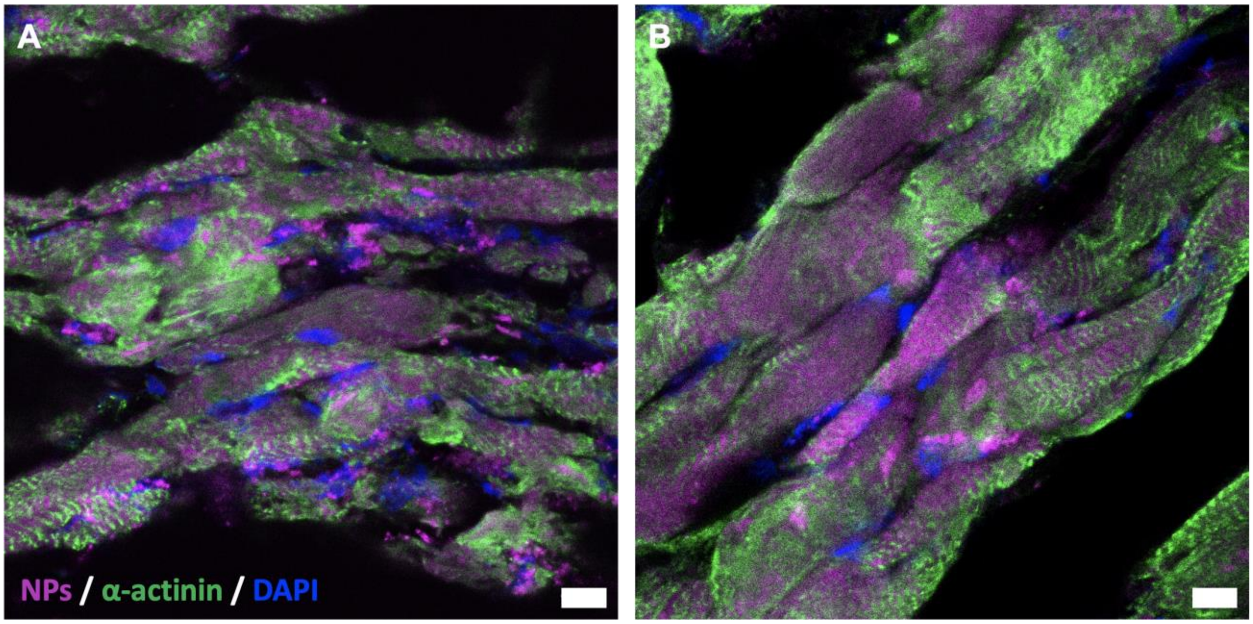

**Figure S12.** DNPs accumulation at 1 day post-injection colocalized strongly with cardiomyocytes in the infarcted region.

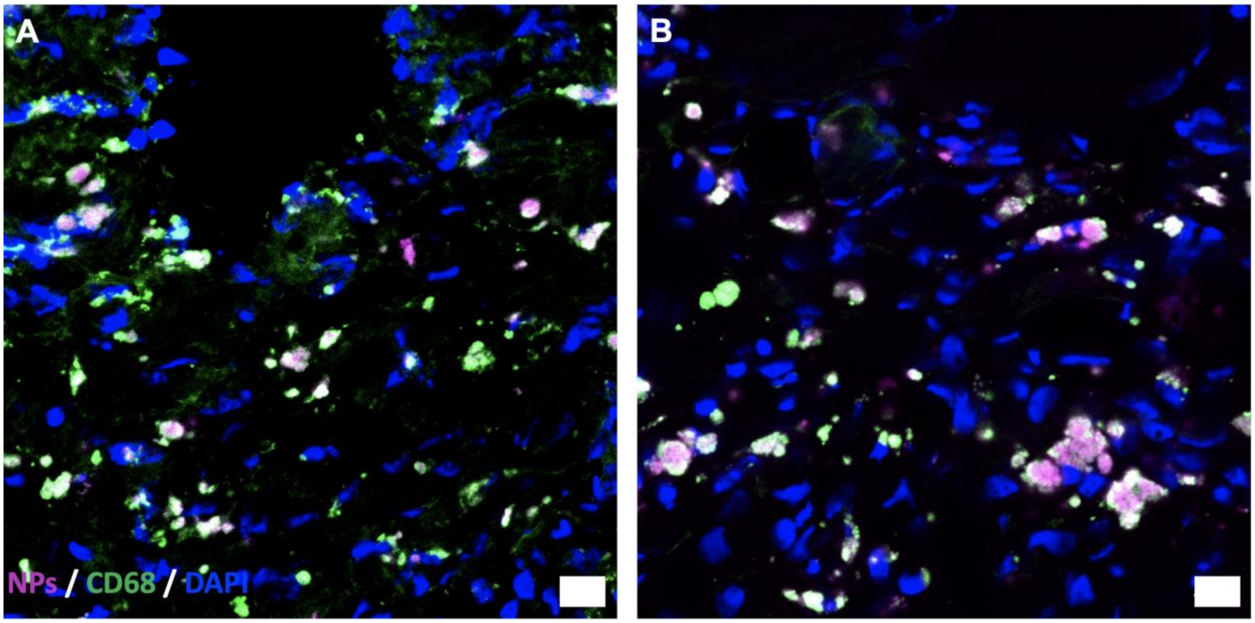

**Figure S13.** At (A) 14 and (B) 28 days post-injection, DNPs colocalized with $CD68^+$ macrophages in the heart.
